# A DNA-based voltmeter for organelles

**DOI:** 10.1101/523019

**Authors:** Anand Saminathan, John Devany, Kavya S. Pillai, Aneesh T. Veetil, Michael Schwake, Yamuna Krishnan

## Abstract

The role of membrane potential in most intracellular organelles remains unexplored because of the lack of suitable probes. We describe a DNA-based fluorescent reporter that quantitates membrane potential and can be targeted to specific organelles in live cells. It is equipped with a voltage sensitive fluorophore, a reference fluorophore for ratiometric quantification, and acts as an endocytic tracer. We could thereby measure the membrane potential of different intracellular organelles in living cells, which has not been possible previously. Our understanding of how membrane potential regulates organelle biology is poised to expand through the use of these new sensors.

**One Sentence Summary:** Using a DNA-based voltmeter we can non-invasively measure the membrane potential of specific organelles in live cells.

## Main Text

Membrane potential is a key property of all biological membranes and underlies how membranes transduce chemical signals. It is therefore a fundamental signaling cue in all cells. (*1*–*5*). Although the membrane potential of the plasma membrane is relatively straightforward to measure due to its accessibility to electrical and chemical probes, that of intracellular organelles is not. The membranes of intracellular organelles enclose lumens of very different ionic compositions from the cytosol, and are therefore expected to maintain distinctive membrane potentials (*6*). In fact, electrophysiology on purified mitochondria and lysosomes separated from disrupted cells have revealed that their membrane potentials are −140 mV (lumen negative) and +100 mV (lumen positive) respectively (*7*, *8*) compared to −70 mV (cytosol negative) for the plasma membrane of a resting cell.

Electrophysiology on isolated organelles indicate that membrane potential could be a major regulator of organelle function (*9*). The mitochondrial membrane potential generated by the Krebs cycle is harnessed to transport ions and charged compounds between the cytosol and the mitochondrial lumen (*10*). Its high negative membrane potential changes in concert with cellular metabolism thereby regulating mitochondrial fission and fusion (*11*, *12*). In fact, depolarized mitochondria cannot undergo effective fusion, and are degraded by mitophagy (*13*).

Lysosomes harbor several voltage-gated ion channels and transporters, the functions of many of which are still unknown (*16*, *17*). Electrophysiology of isolated lysosomes reveal that membrane potential regulates lysosomal functions such as the refilling of its lumenal calcium (*14*) or its fusion with other organelles (*15*).

Importantly, the role of membrane potential in the function of several intracellular organelles cannot be addressed as they are not amenable to electrophysiology and there is a paucity of organelle-specific probes (*8*). Genetically-encodable voltage indicators based on fluorescent proteins are desirable due to their non-invasive nature. However, such probes introduce a capacitive load on the membrane as follows. Generally, they report on membrane potential by repositioning themselves within the membrane depending on their charge state that changes as a function of membrane potential. Thus, the introduction of several probe molecules increases the capacitive load on the membrane, thereby altering the true current versus voltage characteristics of the membrane. Thus, protein-based indicators are incompatible with smaller organelles as their high capacitive loads and overexpression could significantly alter organellar membrane potential (*22*). Further, the pH sensitivity of fluorescent proteins limits their applicability in organelles where lumenal pH and membrane potential are codependent (*21*). In contrast, electrochromic hemicyanine dyes and photoinduced electron transfer (PeT) based voltage sensitive dyes are particularly attractive due to their low capacitive loads, high temporal resolution, photostability and response range (*18*, *19*). However, unlike proteins, voltage sensitive dyes cannot be targeted to specific organelles (*20*).

We have created a DNA-based, organelle-targetable, ratiometric voltage indicator called *Voltair*, with which we can non-invasively measure the membrane potential of specific organelles in live cells. DNA is a versatile, programmable scaffold that is highly suited to chemically imaging living systems (*23*–*27*). A range of detection chemistries displayed on DNA nanodevices have been used to quantitatively map diverse analytes in living systems (*26*–*29*). By leveraging the precise stoichiometry of DNA hybridization, one can incorporate a reference fluorophore and the desired detection chemistry in a well-defined stoichiometry to yield ratiometric probes (*30*). Further, one can display molecular trafficking motifs on DNA nanodevices and localize the latter within sub-cellular organelles (*26*, *31*, *32*).

By targeting *Voltair* to the endo-lysosomal pathway using scavenger receptor-mediated endocytosis, we measured the membrane potential of the early endosome, the late endosome and the lysosome in live cells. We could thereby measure changes in membrane potential that occur as a function of endosomal maturation, which has not been previously possible. We then reprogrammed *Voltair* to engage either the transferrin receptor or furin and thereby localize in either the recycling endosomes or the trans-Golgi network. We could thus measure the membrane potential of the recycling endosome and trans Golgi network in live cells.

Thereafter we demonstrate the capacity of *Voltair* to provide new insights in two contexts. In the first example we measure the contribution of the electrogenic Vacuolar H^+^ - ATPase to the membrane potential of recycling endosomes and trans Golgi network. In the second, we describe a new role for lysosomal membrane potential in transducing how the cell senses nutrient depletion. Using *Voltair* we show that mTORC1 dissociation reduces lysosomal membrane potential by opening the TPC channel. The low lysosomal membrane potential promotes the activity of the voltage-gated K^+^ channel Slo1 which releases lysosomal Ca^2+^ via feedback with the TRPML1 channel to initiate lysosome biogenesis. The ability to measure membrane potential in intact live cells was essential to establish this molecular connectivity.

### Design and Characterization of *Voltair* devices

Our *Voltair* probes comprise a 38-base pair DNA duplex with three functional modules (Fig 1A and Table S1). The first module, denoted D_v_, is a 38-mer single-stranded DNA conjugated at its 3′ end to a previously characterized voltage sensing dye (RVF), using click chemistry described in Materials and Methods (Fig S1-S4) (*33*, *34*). The fluorophore in RVF is quenched by photoinduced electron transfer (PeT) due to hyperconjugation with the lone pair of electrons on its dimethyl aminobenzyl moiety (Fig 1B) (*18*). When a potential difference is applied along the long axis of RVF, the local electric field modulates electron transfer, and therefore quenching, which affects RVF fluorescence (*33*). Specifically, plasma-membrane depolarization decreases electron transfer and increases fluorescence compared to that at resting potential (Fig 1A, Movie S1). RVF has high photostability, no capacitive load, pH insensitivity from pH 4.5 – 7.5, which makes it suitable to interrogate diverse membranes at different physiological pH (Fig S5) (*35*, *36*).

**Fig. 1.**
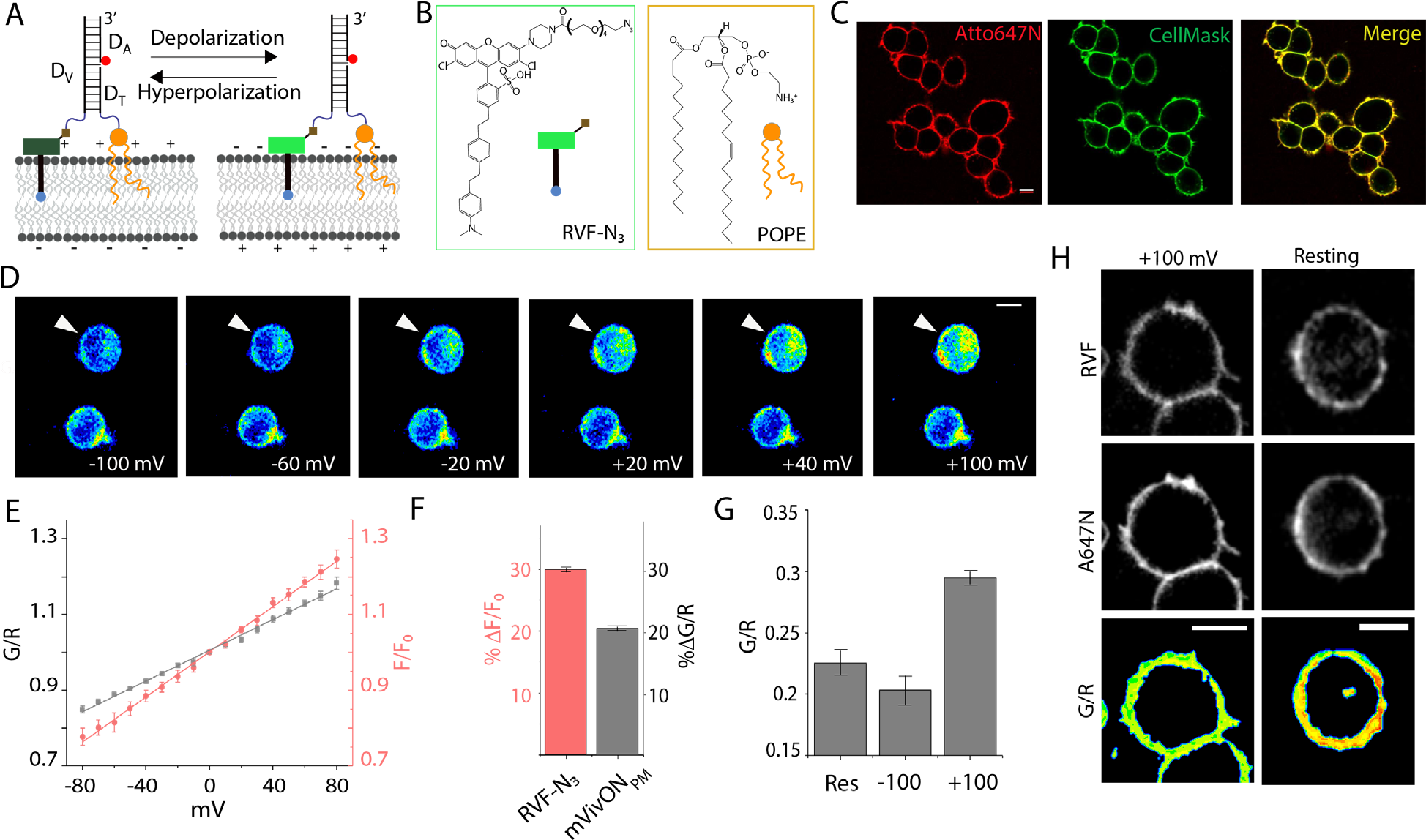
Design and characterization of *Voltair* probes. (A) Schematic of the working principle of *Voltair*^*PM*^: A voltage sensitive dye (RVF) is conjugated to a DNA duplex that is membrane-tethered by attachment to a lipid anchor (POPE). The DNA duplex has a reference dye (Atto647N, red sphere) that together with RVF reports membrane potential ratiometrically. (B) Structure of a conjugatable version of RVF (RVF-N_3_) and the lipid anchor POPE. (C) Colocalization of *Voltair*^*PM*^ with plasma membrane marker CellMask^TM^ orange in HEK 293T cells. (D) Pseudo-color images show pixel-wise ratio of RVF to Atto647N fluorescence intensities (G/R) as a function of membrane potential applied by whole cell voltage clamping (white arrow head indicates clamped cell). (E) Sensor response of *Voltair*^*PM*^ (gray trace) and unmodified RVF (red trace) as a function of plasma membrane potential in HEK 293T cells. G/R values are normalized to that at 0 mV. (F) Percentage change in signal of *Voltair*^*PM*^ (black) and unmodified RVF (red) from 0 to 100 mV. (G, H) G/R values of the plasma membranes of resting cells (Res) and when they are voltage clamped at −100 mV and +100 mV. Data represent the mean ± s.e.m. of three independent trials. Error bar represents the standard deviation for n = 10 cells. Scale bar = 10 μm.

The second module in *Voltair* probe is a reference dye, that is insensitive to membrane potential. Including the reference dye corrects for intensity changes due to different sensor concentrations arising from non-uniform cellular uptake. Thus, ratiometric measures of RVF and reference dye intensities are proportional only to membrane potential. The reference dye, Atto647N, is attached to the 5′ end of D_A_ and is chosen for its high photostability, negligible spectral overlap with RVF, insensitivity to pH, voltage and other ions (red circle, Fig 1A).

The third module in *Voltair* probe variants comprises a targeting moiety to efficiently localize the probe, either at the plasma membrane, or in intracellular membranes of specific organelles. The *Voltair*^*PM*^ variant has a targeting motif that localizes *Voltair* to the plasma-membrane by chemically conjugating the 5′ end of D_T_ to 1-palmitoyl-2-oleoyl-sn-glycero-3-phosphoethanolamine (POPE) moiety via a tetra-ethylene glycol linker (*37*). When *Voltair*^*PM*^ is added to the culture media, the POPE moiety inserts into the outer leaflet of the plasma membrane, anchoring *Voltair*^*PM*^ to the cell surface (Fig 1C, S6). Thus, RVF is inserted in a defined orientation with respect to the membrane potential vector, and the anionic DNA duplex prevents any potential flipping of the RVF moiety in the membrane.

In order to target *Voltair* to intracellular membranes, we used the *Voltair*^*IM*^ variant which leverages the inherent ability of duplex DNA to serve as a ligand for scavenger receptors (*38*). The DNA-based targeting motif imposes an over-riding trafficking signal that enables probe internalization into a specific intracellular organelle. The DNA duplex binds scavenger receptors at the plasma-membrane and undergoes scavenger receptor-mediated endocytosis, trafficking to early endosomes, which mature to late endosomes, and finally to lysosomes, in a time-dependent manner (*25*). Both *Voltair* probes were assembled by annealing equimolar amounts of the component strands to yield probes with a precise 1:1 stoichiometry of the sensing dye (RVF) and reference dye (Atto647N). The formation and integrity of *Voltair*^*PM*^ and *Voltair*^*IM*^ were confirmed using gel electrophoresis, which indicated >99% yield (Fig S3). Both *Voltair*^*PM*^ and *Voltair*^*IM*^ report on membrane potential by sequentially exciting RVF and Atto647N and monitoring their emission intensities at λ_em_ = 540 nm (G for RVF) and λ_em_ = 665 nm (R for Atto647N) (Fig S4). We describe later in detail how the targeting motifs work to successfully localize the *Voltair* probes selectively at the plasma membrane or inside an organelle.

We first characterized the voltage-sensitive response of *Voltair*^*PM*^ using whole-cell voltage clamping using the set-up described in Fig S7 (methods) (*39*). The plasma membrane of HEK 293T cells was efficiently labeled by *Voltair^PM^*, as seen by colocalization with the membrane stain Cellmask_TM_ (Fig. 1C, S6). Next, a single *Voltair*^*PM*^ labeled cell was voltage clamped from −100 mV to +100 mV, at increments of 10 mV, and fluorescence images of RVF (G channel) and Atto647N (R channel) were sequentially acquired at each clamped voltage, from which the pseudo-colored G/R images were generated as described (Fig 1D, methods). Notably, the whole-cell intensity of images in the G channel changed as a function of the applied voltage, while the R channel was constant (Fig S8, Movie S2). Unclamped cells showed negligible variation in either channel. The uniform 1:1 ratio of RVF: Atto647N in *Voltair*^*PM*^ yield highly reproducible G/R values as a function of applied voltage as seen in the calibration plot of *Voltair*^*PM*^ (Fig 1E). The fold change in G/R signal from 0 mV to +100 mV (ΔG/R) was found to be 1.2 for *Voltair*^*PM*^ which matched well with that of unmodified RVF (*33*) (Fig 1F). Based on the G/R value obtained for resting cells labeled with *Voltair^PM^*, we found the resting potential to be −50 mV using the calibration profile, consistent with previous measurements (*40*) (Fig 1G, H).

### Targeting *Voltair*^*IM*^ to membranes of specific endocytic organelles

Given the favorable reporter characteristics of *Voltair*^*PM*^ we then localized the variant, *Voltair*^*IM*^, in organelles along the endo-lysosomal pathway using scavenger receptor mediated endocytosis (Fig 2A). HEK 293T cells do not express scavenger receptors (*41*). While this is ideal for plasma membrane immobilization of *Voltair*^*PM*^, it disfavors endosomal uptake of *Voltair* probes. Therefore, we over-expressed human macrophage scavenger receptor (hMSR1) fused to CFP in HEK 293T cells by transient transfection. Transfected cells effectively endocytosed *Voltair*^*IM*^ through receptor-mediated endocytosis (Fig 2B-C). Colocalization between hMSR1-CFP and *Voltair*^*IM*^ revealed that its uptake occurred through the scavenger receptor pathway (Fig 2B). This was reaffirmed by competing out *Voltair*^*IM*^ uptake using excess maleylated BSA, a good ligand for scavenger receptors (Fig 2C, S9) (*42*).

**Fig. 2.**
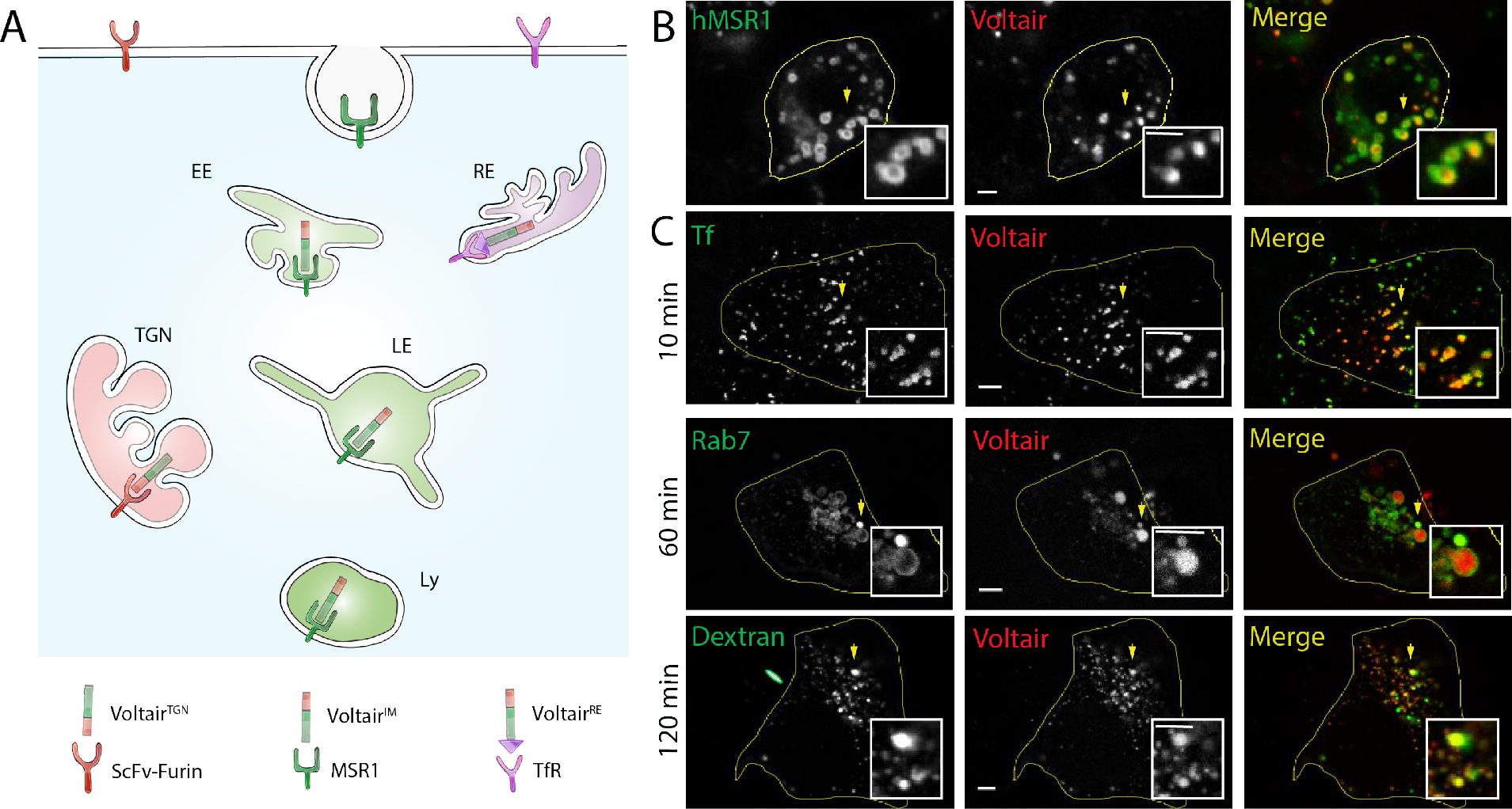
Targeting *Voltair*^*IM*^ to membranes of specific endocytic organelles. (A) Schematic of targeting strategy: *Voltair*^*IM*^ undergoes scavenger receptor mediated endocytosis by binding scavenger receptors. Endocytosed *Voltair*^*IM*^ traffics in a time-dependent manner from the plasma membrane to the early endosome, the late endosome and then the lysosome. (B) Colocalization between internalized *Voltair*^*IM*^ (red channel) and human macrophage scavenger receptors (hMSR1-CFP, green channel) transfected in HEK 293T cells. (C) Co-localization between various endocytic organelle markers (green channel) and *Voltair*^*IM*^ (red channel) at the indicated chase times. Early endosomes are labelled with Transferrin-Alexa546 (Tf), late endosomes are labelled with Rab7-mRFP (Rab7) and lysosomes are labelled with TMR dextran (Dextrans). Scale bar = 5 μm.

We then determined the timepoints of localization of internalized *Voltair*^*IM*^ at each stage along the endo-lysosomal pathway in HEK 293T cells (Fig 2D-E). For this, we performed time-dependent colocalization experiments with various endocytic markers. Briefly, HEK 293T cells expressing hMSR1 were pre-labeled with endocytic markers i.e., Alexa546-labeled transferrin, Rab7-mRFP and TMR-Dextran, that labeled early endosomes, late endosomes or lysosomes, respectively (*43*, *44*). These cells were then pulsed with singly-labeled *Voltair*^*IM*^. Colocalization was monitored as a function of different chase times (Fig 2D-E, S10). We found that *Voltair*^*IM*^ localized in early endosomes at ~10 min, in late endosomes at ~60 min, and in lysosomes at ~120 min. In *Voltair*^*IM*^, the RVF moiety functions both as a voltage sensitive dye as well as a lipid anchor that tethers *Voltair*^*IM*^ to the lumenal face of the organelle membrane (Fig S11). In parallel, the duplex DNA moiety allows scavenger receptor binding and also allows RVF insertion into the intracellular membrane surrounding the receptor, and therefore overrides the affinity of RVF to the plasma membrane (Fig S12). Thus, integration onto a duplex DNA scaffold successfully imposes the scavenger receptor-mediated endocytic program on to RVF.

### Calibration of *Voltair* probes in intracellular organelles

Next, we mapped the response characteristics of *Voltair* in intracellular membranes. To achieve this, we developed a method to substitute the cytosol with buffer of a desired composition without affecting the integrity of fluorescently labeled lysosomes (Fig S13, S14). This uses digitonin treatment of labeled cells in the presence of a buffer of desired ion concentrations. Digitonin treatment creates 30 nm wide pores on the plasma membrane through which small molecules such as ATP and ion content of the cytosol and the external buffer rapidly equilibrate (Fig 3A) (*45*). Further, lysosomal pumps and transporters are functional, lysosomal membrane integrity is preserved thus enabling the active maintenance of lysosomal membrane potential in these digitonin treated cells.

**Fig. 3.**
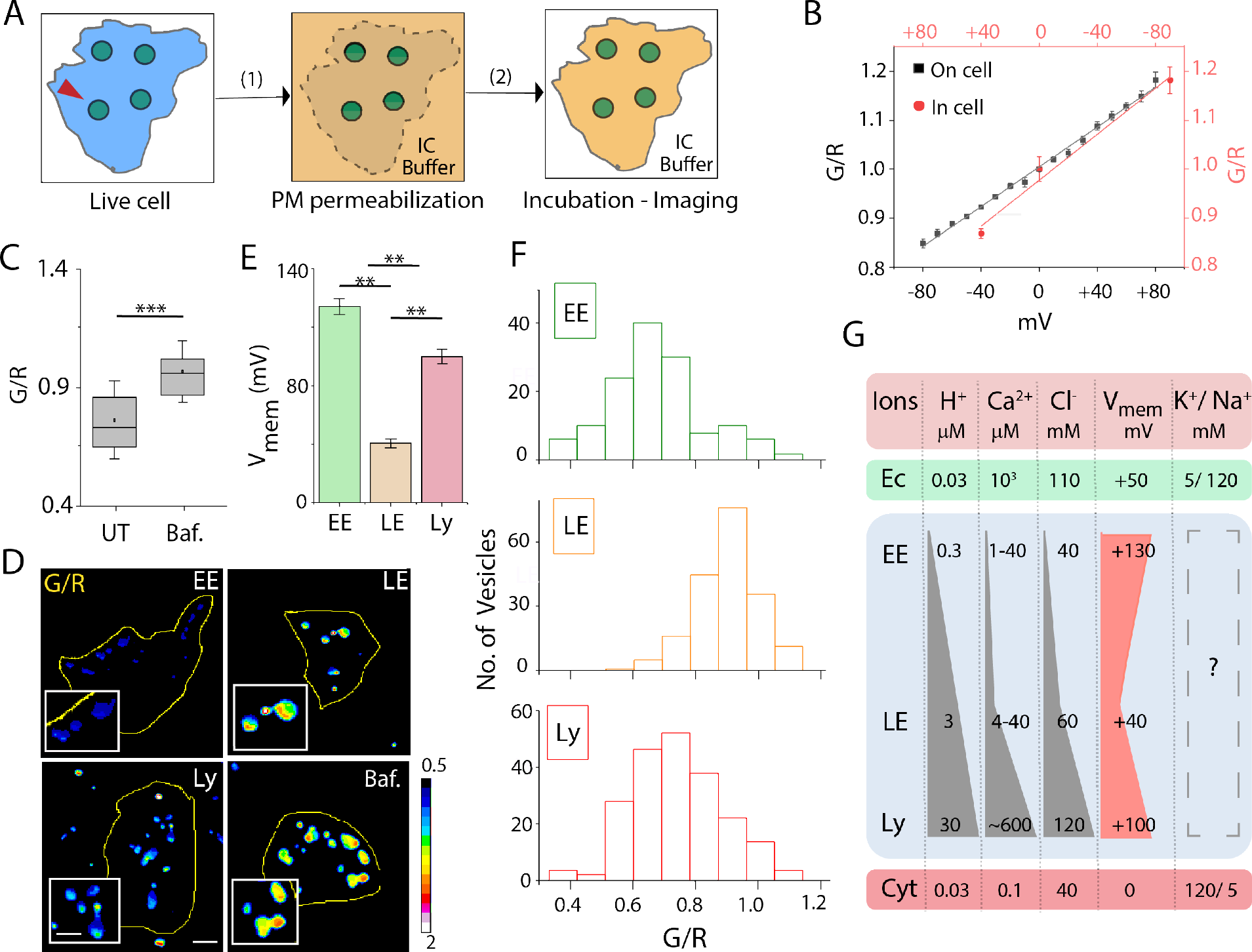
*Voltair*^*IM*^ measures membrane potential of intracellular organelles. (A) Schematic of substituting the cytosol with a desired buffer of ions and small molecules (IC buffer). Red arrowheads indicate *Voltair*^*IM*^ labeled lysosomes. (1) Cells are permeabilized with 10 μm digitonin in IC buffer for 2 minutes (2) Cells are washed and incubated in IC buffer and then imaged. (B) G/R values of *Voltair*^*IM*^ (red) labeled lysosomes and *Voltair*^*PM*^ (black) labeled plasma membranes of HEK 293T cells clamped at specific lysosomal and plasma membrane potentials respectively. Error bars indicate mean ± s.e.m. of three independent measurements. (C) G/R values of *Voltair*^*PM*^ labelled lysosomes in presence (Baf) and absence (UT) of 1 μM Bafilomycin. Error bars indicate mean ± s.e.m. of three independent measurements; \*\*\**P* < 0.001 (Unpaired two sample t test). (D) Representative pseudocolor G/R images of the indicated organelles labeled with *Voltair*^*IM*^. Also shown is the corresponding image of *Voltair*^*IM*^ labeled lysosomes treated with bafilomycin (1μM). (E) Measured membrane potentials of early endosomes (EE), late endosomes (LE) and lysosomes (Ly). Error bars indicate mean ± s.e.m. of six independent measurements; \*\**P* < 0.01 (one-way ANOVA with Tukey *post hoc* test). (F) Histograms showing the spread of G/R values of n =200 organelles at each stage. (G) Cytosolic (Cyt), extracellular (Ec) and organellar concentrations of main ions that maintain membrane potential. Scale bar = 5 μm.

We constructed an in-cell calibration plot of membrane potential which reveals that reporter characteristics of *Voltair*^*IM*^ in intracellular membranes quantitatively recapitulates that of *Voltair*^*PM*^ at the plasma membrane (Fig 3B). To achieve this, we labeled lysosomes with *Voltair*^*IM*^ and, using digitonin treatment, substituted the cytosol with buffers containing specific ionophores that have previously been shown to create well-defined values of membrane potential in isolated lysosomes. Valinomycin (50 μM) results in a membrane potential of +40 mV considering the lumen to be positive (*46*), a cocktail of valinomycin and monensin (50 μM) with 150 mM K^+^ neutralizes lysosomal membrane potential (*47*), while FCCP (50 μM) in the absence of ATP creates a membrane potential of −90 mV across the lysosomal membrane (*48*). G/R values for ~50 *Voltair*^*IM*^ labeled lysosomes for each of these conditions were plotted as a function of the expected lysosomal membrane potential (Fig. 3B, S15). *Voltair*^*IM*^ shows 25%-fold change in G/R signal per 100 mV, which is highly consistent with the reporter response of *Voltair*^*PM*^ (Fig. 3B). We use the calibration profile of *Voltair*^*IM*^ to compute the membrane potential of any intracellular organelle.

We then confirmed that this intracellular calibration plot was indeed reliable, by measuring lysosome membrane potential *in situ* before and after inhibiting V-ATPases and comparing our results with previous measurements on isolated organelles from disrupted cells. The electrogenic proton pump V-ATPase hydrolyses ATP to acidify organelles and thereby generates membrane potential across organelle membrane (*49*). Inhibiting V-ATPase with bafilomycin stops proton transport and this is theoretically expected to neutralize lysosomal membrane potential (*49*, *50*). This has indeed been observed in the case of purified synaptic vesicles (*51*). We found that *Voltair*^*IM*^ labeled lysosomes showed a large increase in G/R values in the presence of bafilomycin (1 μM) as compared to that in untreated cells, revealing that lysosomal membrane potential was indeed dissipated by inhibiting V-ATPase (Fig 3C-D). From the in-cell calibration plot (Fig 3B), this change corresponded to a difference of −90 mV from the resting membrane potential and is highly consistent with other studies on V-ATPase regulated synaptic vesicles (*51*). (Fig 3C-D).

### Membrane potential as a function of endosomal maturation

We then measured the resting membrane potential of endocytic organelles in intact, live cells as a function of endosomal maturation. Early endosomes, late endosomes and lysosomes were each specifically labelled with *Voltair*^*IM*^, as described earlier in Figure 2D (Fig S16). Fluorescence images in the G and R channels were acquired and the G/R values of ~200 organelles were computed for each endosomal stage (Fig 3D-E, S17). Figure 3F shows the distribution of G/R values at each stage along the endocytic pathway and from the intracellular calibration plot we could determine the corresponding organelle membrane potential (Fig 3E, S17, S18) Considering the lumen to be positive and the cytoplasmic face as negative, the membrane potential of lysosomes (V_Ly_) was found to be +100 mV. This is in line with other electrophysiological studies of lysosomal membrane potential in different cell types that were found to range between +30 mV to +110 mV (*8*). However, no such measurements exist for early or late endosomes. Measurements with *Voltair*^*IM*^ revealed the membrane potential of early endosomes (V_EE_) and late endosomes (V_LE_) to be +130 mV and +40 mV respectively (Table S2).

Surprisingly the gradient of membrane potential accompanying endosomal maturation did not reflect that of many known ions (Fig 3G). The concentrations of protons, chloride and calcium increase progressively during endosomal maturation (Fig 3G). In contrast, membrane potential is highest in the early endosome, drops ~3 fold in the late endosome and increases again in lysosomes. With respect to cytoplasmic concentrations, the levels of lumenal [Ca^2+^] and [Cl^−^] for the early endosome or the lysosome, are very similar i.e., ~10^2^-10^3^ fold higher [Ca^2+^], 1-2-fold higher [Cl^−^] (Fig 3G) (*26*, *52*). However, the lysosome lumen has ~10^3^ fold higher [H^+^], while the early endosome lumen has only ~10 fold higher [H^+^] than the cytosol(*25*). Yet the observed membrane potentials for both these organelles are consistent with the lumenal face of the lysosome being less positive, rather than more positive, compared to the early endosome (V_EE_ = +130 mV; V_LY_ = +100 mV). Considering the low abundances of other free ions, this provides a compelling case for Na^+^ and K^+^ transporters or exchangers in maintaining the high positive membrane potential of the early endosome. In fact, late endosomes are posited to contain high K^+^, which is consistent with our observations (*53*). Our hypothesis is supported by observations at the plasma membrane. The differences in [Ca^2+^] and [Cl^−^] across the plasma membrane are comparable, yet its membrane potential is set by the differences in [Na^+^] and [K^+^] across the membrane (*54*, *55*).

### Membrane potential of organelles along recycling and retrograde trafficking pathway

We then showed that *Voltair* reporters can interrogate organelles beyond the endolysosomal pathway. We re-designed *Voltair* to give *Voltair*^*RE*^ and *Voltair*^*TGN*^ that could access the recycling and retrograde pathways and measure the membrane potential of the recycling endosome and the trans-Golgi network respectively. These organelles have been only hypothesized to have membrane potential which is thought to be regulated by the electrogenic activity of vacuolar H^+^-ATPases. *Voltair*^*RE*^ displays an RNA aptamer against the human transferrin receptor (Fig S20) (*26*) and was uptaken by the transferrin receptor pathway. *Voltair*^*RE*^ was trafficked to recycling endosomes and evidenced by co-localization with Alexa 546 labelled transferrin (Fig 4A, B). *Voltair*^*TGN*^ was targeted to the trans-Golgi network in HEK 293 cells expressing furin fused to a single-chain variable fragment recombinant antibody, ScFv (*24*, *31*). The ScFv domain selectively binds any DNA duplex containing a d(AT)_4_ sequence. *Voltair*^*TGN*^ included a d(AT)_4_ sequence in the duplex so that the ScFv-Furin chimera acts as a carrier protein for the former. *Voltair*^*TGN*^ is thus trafficked along the retrograde pathway localizing in the trans Golgi network evidenced by clear colocalization with TGN46-mCherry and absence of colocalization with endocytic organelles (Fig 4C, D, S21).

**Fig. 4.**
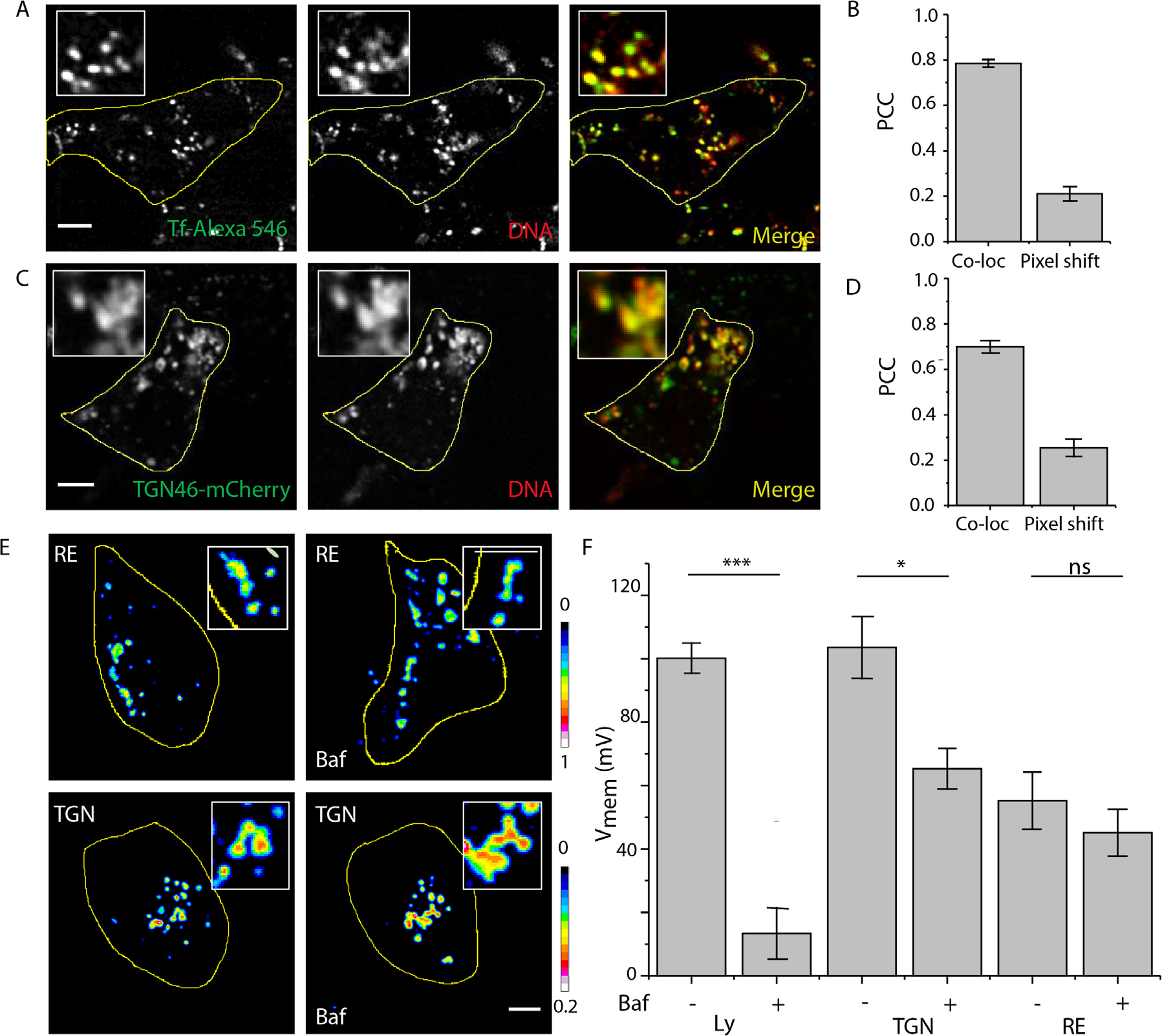
Membrane potential of organelles along recycling and retrograde trafficking pathway. (A) Colocalization between internalized *Voltair*^*RE*^ (red channel) and recycling endosome marker (Transferrin-Alexa546, green channel) in HEK 293T cells. (B) Pearson’s correlation coefficient (PCC) of colocalization in (A). (C) Colocalization between internalized *Voltair*^*TGN*^ (red channel) and trans-Golgi network marker (TGN46-mCherry, green channel) in ScFv-Furin transfected HEK 293T cells. Scale = 5 μm, inset = 5 μm. (D) Pearson’s correlation coefficient (PCC) of colocalization in (C). (E) Representative pseudo-color G/R images of RE and TGN of HEK 293T cells in absence and presence of bafilomycin (500 nM). Scale = 5 μm. (F) Resting membrane potential of organelles and changes upon inhibition of V-ATPase. Error bars indicate mean ± s. e. m; ns, not significant (*P* > 0.05); \*\*\**P* < 0.001, \**P* < 0.05 (one-way ANOVA with Tukey *post hoc* test).

We then measured the resting membrane potential of the recycling endosome and trans-Golgi network, neither of which has been previously possible, and evaluated the contribution of V-ATPase to membrane potential in each organelle. *Voltair*^*RE*^ labelled cells were imaged in the G and R channels, and G/R values of ~50 recycling endosomes were computed (Fig 4E, F). We found that the membrane potential of recycling endosome (V_RE_) was +55±11 mV (lumen positive), similar to plasma membrane (cytosol negative). When V-ATPase activity was inhibited by treating cells with 500 nM bafilomycin, the membrane potential V_RE_ showed no change revealing a negligible contribution of V-ATPase in these membranes(*56*). The magnitude and the V-ATPase dependence of membrane potential in the recycling endosome mirrors that of the plasma membrane, indicating that both membranes share similar electrical characteristics.

*Voltair*^*TGN*^ labelled cells were similarly imaged and G/R values of the TGN in ~30 cells was computed, we found that the membrane potential of the trans Golgi network (V_TGN_) was +103±16 mV (lumen positive) (Fig 4E, F). The high membrane potential of the trans-Golgi network is surprising as the Golgi has been envisaged to have negligible membrane potential, since its high permeability to K^+^ ions could electrically balance the H^+^ influx needed to acidify its lumen (*57*). In contrast to what we observed for recycling endosomes, V-ATPase inhibition substantially reduced V_TGN_. However, unlike in lysosomes, the membrane potential of the TGN could not be completely neutralized and still showed +65 ± 10 mV across the membrane. This suggests that other electrogenic transporters at the TGN, possibly Na^+^/K^+^ ATPases could significantly contribute to the V_TGN_ (*58*–*60*).

### mTORC1 recruitment to the lysosome alters lysosomal membrane potential

We then explored the capability of our probe technology to deliver new insights into more well-investigated problems. The absence of nutrients in the cell is sensing by mammalian target of rapamycin complex (mTORC1) to initiate lysosome biogenesis and upregulate autophagy (*61*). Normally, mTORC1 is present on lysosomes where it regulates the activity of diverse lysosomal proteins (*62*). When nutrient levels plummet mTORC1 dissociates from the lysosome which triggers the release of lysosomal Ca^2+^ through the TRPML1 channel and initiate lysosomal biogenesis (*63*). How mTORC1 dissociation actually activates the TRPML1 channel is not yet understood (*64*). We show a new role for lysosomal membrane potential, where it transduces mTORC1 dissociation to TRPML1 through the voltage dependent channel, Slo1 (Fig 5A).

**Fig. 5.**
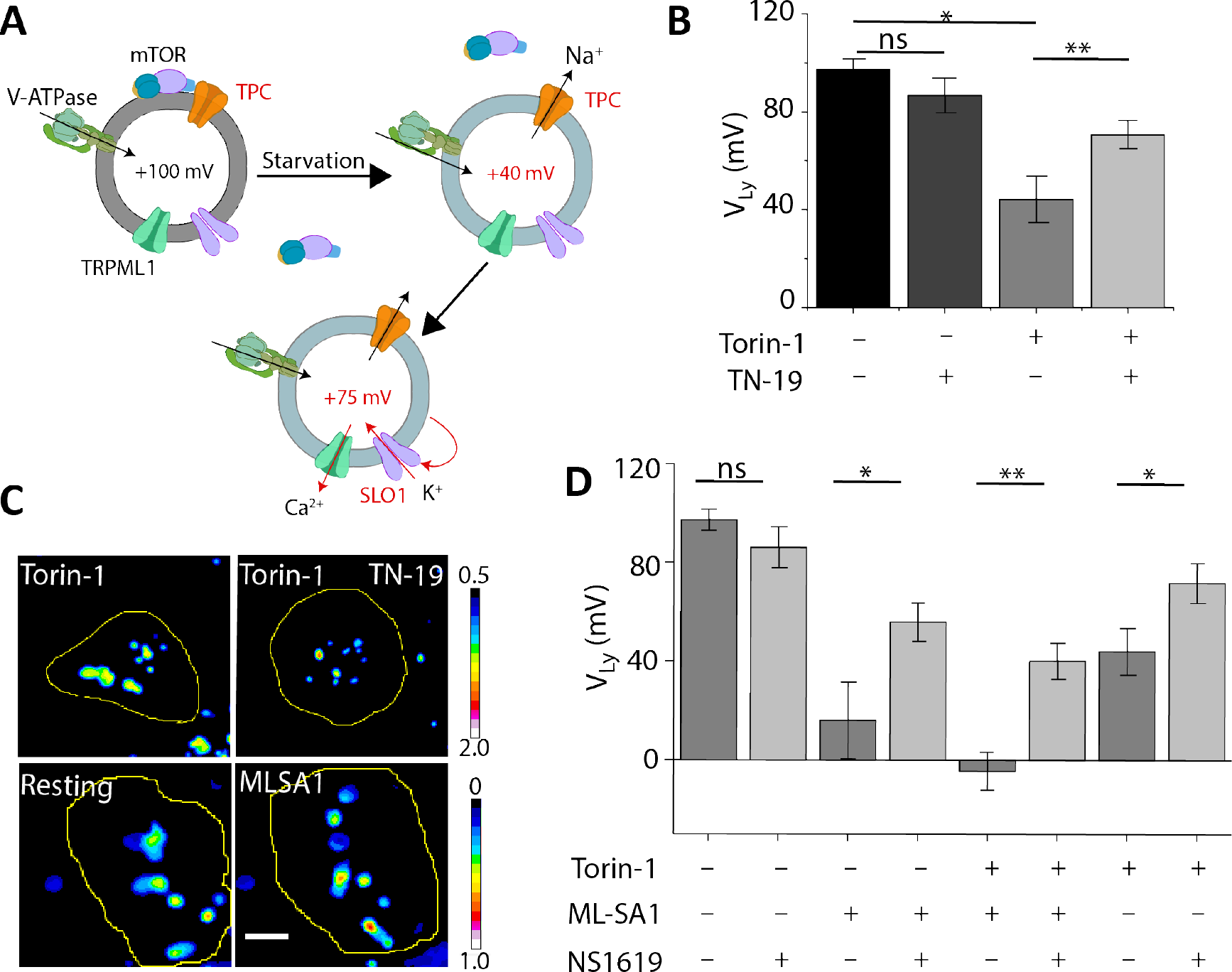
Dissociation of mTORC1 from the lysosome reduces the membrane potential and activates Slo1. (A) When mTORC1 dissociates from the lysosome the TPC channel opens, releasing Na^+^ and depolarizing the lysosome to +40 mV. Low membrane potential promotes the opening of the voltage gated channel Slo1, which, by coupling to TRPML1 releases lysosomal calcium. (B) Lysosomal membrane potential changes in the presence of the mTORC1 inhibitor Torin-1 (1μM) and the TPC channel inhibitor TN-19 (1μM). Error bars indicate mean ± s.e.m. of three independent measurements; ns, not significant (*P* > 0.05); \*\**P* < 0.01 (one-way ANOVA with Tukey *post hoc* test). (C) Representative pseudo-color G/R images of lysosomes in HEK 293T cells in presence of indicated pharmacological agents (1μM). Scale = 5μm. (D) Lysosomal membrane potential in presence of Torin-1 (1 μM), the TRPML-1 activator ML-SA1 (20 μM), and the Slo1 activator, NS1619 (15 μM). Error bars indicate mean ± s.e.m. of three independent experiments; ns, not significant (*P* > 0.05); \*\**P* < 0.01; \**P* < 0.05 (Unpaired two sample t test).

While on the lysosome, mTORC1 inhibits the Two Pore Channel (TPC) (*65*). Although TPC channel activation has been shown to depolarize the lysosome(*15*, *66*) there is no direct evidence that mTORC1 dissociation could cause lysosome depolarization in intact cells. We therefore triggered mTORC1 dissociation by inhibiting mTORC1 with Torin-1 (*67*). Torin-1 (1 μM) treatment dramatically reduced lysosome membrane potential (V_Ly_)from +100 mV to +40 mV (Fig. 5 B, C). When the experiment was repeated in the presence of the TPC channel inhibitor *trans*-ned-19 (TN-19, 1 μM) (*68*), we observed only a minor reduction in V_Ly_ from +100 mV to +75 mV, indicating that indeed lysosome depolarization due to mTORC1 dissociation in cells is mediated mainly by the TPC channel.

We then considered whether the extensive depolarization of the lysosome caused by mTORC1 dissociation could affect a voltage-gated channel downstream. Slo1 is a lysosomal BK channel, responsible for K^+^ influx into the lysosome, which, if activated would make the lumenal face of the lysosomal membrane more positive and increase V_Ly_ (*69*). In parallel, studies on the plasma membrane BK channel indicate that the cytosolic Ca^2+^ binding domain and voltage sensitive S4 domain respond to high cytosolic Ca^2+^ and low membrane potential to drive K^+^ transport (*70*). Thus, we posited that lysosome depolarization could modulate Slo1 activity that, could drive lysosomal Ca^2+^ release by feedback with the TRPML1 channel (*14*, *69*).

By measuring membrane potential in the presence of chemical modulators of Slo1 and TRPML1 channels, we find that lysosome depolarization promotes Slo1 activity which triggers TRPML1 channel opening. We used NS1619, which is a small molecule that opens Slo1 by left-shifting its half-activation voltage (V_0.5_) (*68*, *71*). In other words, NS1619 causes Slo1 to open only at lower V_Ly_ (*14*). In the presence or absence of NS1619, *Voltair*^*IM*^ labeled lysosomes V_Ly_ remained invariant. This is expected, because the large positive V_Ly_ values of mTORC1-associated lysosomes should render NS1619 ineffective at opening Slo1. Thus, in mTORC1-associated lysosomes Slo1 is inactive and likely requires a considerable drop in V_Ly_ and/or elevation in cytosolic Ca^2+^ to open.

We therefore elevated cytosolic Ca^2+^ in the vicinity of Slo1 by adding ML-SA1, a TRPML1 channel activator that releases lysosomal Ca^2+^ (Fig 4D) (*72*). We found that this lowers V_Ly_ to +20 mV. We then checked whether these depolarized conditions promoted the opening of Slo1 by adding NS1619, which if effective, would be expected drive V_Ly_ to more positive values. Interestingly, treating cells with NS1619 in the presence of ML-SA1 indeed shifted V_Ly_ to +60 mV. This revealed that Slo1 was clearly now responsive to NS1619, as it drove V_Ly_ up by +40 mV. Thus, opening the TRPML1 channel releases lysosomal Ca^2+^ depolarizing the lysosome and promoting the activity of Slo1 (Fig 5D).

By analogy we considered whether opening the TPC channel, which also depolarizes the lysosome by releasing lysosomal Na^+^, could similarly activate Slo1. Importantly, when cells were treated with both Torin-1 and NS1619, *Voltair*^*IM*^ labeled lysosomes showed that V_Ly_ was driven up to +75 mV, i.e., an increase of 35 mV in the positive direction. This indicated that NS1619 was highly effective at opening Slo1. Treating cells with ML-SA1 and Torin-1 dramatically depolarized lysosomes, with V_Ly_ values reaching as low as −5 mV. Again, addition of NS1619 to the cocktail repolarized lysosomes to V_Ly_ +40 mV, i.e., an increase of 45 mV in the positive direction. This confirms that lysosomes mTORC-1 dissociation depolarizes lysosomes, and the low membrane potential causes the voltage-dependent channel Slo1 to open. Then Slo-1, through feedback, causes the TRPML1 channel to release lysosomal Ca^2+^ and initiate lysosome biogenesis.

## Summary

Protein-based voltage indicators while powerful, have limited applicability to organelles, due to their pH sensitivity and capacitive load, as organelles are both small and acidic. Voltage sensitive dyes have low capacitive loads and are also pH insensitive, but, on their own, cannot be targeted to organelles. In contrast, *Voltair* probes succeed because they unite the advantages of voltage sensitive dyes with the organelle-targetability of proteins to non-invasively measure the membrane potential of organelles. These reporters leverage the 1:1 stoichiometry of DNA hybridization to integrate the following functions with stoichiometric precision: (i) a voltage sensing function using voltage sensitive dyes (ii) an internal reference dye for ratiometric quantitation and (iii) an organelle targeting function for stable organelle localization.

Using *Voltair* probes we provide direct evidence that when mTORC1 dissociates from the lysosome, it reduces the membrane potential of the lysosome by relieving its inhibition of the TPC channel. The depolarized the lysosomal membrane promotes the opening of the voltage gated Slo1 channel that, through well-documented feedback with the TRPML1 channel, facilitates the release of lysosomal Ca^2+^ (*14*, *63*).

mTORC1 inhibition sequesters essential amino acids within the lysosome in a highly coordinated manner by impeding their transport across the lysosomal membrane (*73*). Conversely, while resident on the lysosome, mTORC1 facilitates the simultaneous transport of several amino acids from the lysosome into the cytosol (*74*). Thus, mTORC1 has been postulated to modulate the overall nature of the lysosome, adapting it from degrade-and-recycle mode to storage mode (*73*). Such adaptation would require mTORC1 to simultaneously modulate multiple transporters and channels on the lysosome. This can be now envisaged through the ability of mTORC1 to alter lysosomal membrane potential, given that our in-cell membrane potential measurements reveal a change large enough to modulate the function of voltage gated ion channels such as Slo1.

*Voltair* probes combine the attractive reporter characteristics of small molecule voltage sensitive dyes along with the organelle targetability of endocytic tracers. This hybrid technology now enables non-invasive, *in situ* mapping of membrane potential for a range of sub-cellular organelles. For many of these organelles, the endogenous membrane potential was unknown. We have now mapped membrane potential changes that occur during endosomal maturation by addressing organelles on the endolysosomal pathway. Our studies reveal that the early endosome and the lysosome have membrane potentials (lumen positive) as high as +134 ± 17 mV and +100 ± 13 mV, while the late endosome shows a much lower membrane potential of +40 ± 5 mV. These measurements provide compelling evidence for either Na^+^ or K^+^ in establishing the high membrane potential of the early endosome.

Recycling endosomes showed a high degree of similarity with the plasma membrane in terms of the magnitude of the membrane potential as well as its dependence on V-ATPase. The membrane potential of the trans Golgi network, previously hypothesized to be negligible, is revealed to be as high as the lysosome. Yet, unlike the lysosome, it is not completely driven by V-ATPase. Non-invasively interrogating membrane potential in organelles that have proved previously impossible to address, offers the capacity to uncover how these organelles exploit membrane potential to regulate their function.

## Acknowledgments

We thank Professors John Kuriyan and Francisco Bezanilla for critical comments; Kasturi Chakraborty, Joao L. C. Souza, Jary Delgado and Yuanwen Jiang for discussions; the Integrated Light Microscopy and mass spectrometry facilities at the University of Chicago.

## Author contributions

A.S., A.T.V and Y.K. designed the project. A.S., K.S.P., A.T.V. and J.D. performed experiments. M.S. provided a reagent. A.S. and Y.K. analyzed and interpreted the data. A.S. and Y.K. wrote the paper. All authors gave input on the manuscript.

## Data and materials availability

All data is available in the main text or the supplementary materials.

## References and Notes

1. L. F. Santana, E. P. Cheng, W. J. Lederer, How does the shape of the cardiac action potential control calcium signaling and contraction in the heart? J. Mol. Cell Cardiol. 49, 901–903 (2010).

2. F. Helmchen, K. Imoto, B. Sakmann, Ca2+ buffering and action potential-evoked Ca2+ signaling in dendrites of pyramidal neurons. Biophys. J. 70, 1069–1081 (1996).

3. S. A. R. Mousavi, A. Chauvin, F. Pascaud, S. Kellenberger, E. E. Farmer, GLUTAMATE RECEPTOR-LIKE genes mediate leaf-to-leaf wound signalling. Nature. 500, 422–426 (2013).

4. A. Prindle et al., Ion channels enable electrical communication in bacterial communities. Nature. 527, 59–63 (2015).

5. B. Alberts et al., Ion Channels and the Electrical Properties of Membranes (2002).

6. M. Grabe, G. Oster, Regulation of organelle acidity. J. Gen. Physiol. 117, 329–344 (2001).

7. D. G. Nicholls, M. W. Ward, Mitochondrial membrane potential and neuronal glutamate excitotoxicity: mortality and millivolts. Trends Neurosci. 23, 166–174 (2000).

8. X.-P. Dong, X. Wang, H. Xu, TRP channels of intracellular membranes. J. Neurochem. 113, 313–328 (2010).

9. X. Z. Zhong, X.-P. Dong, Lysosome electrophysiology. Methods Cell Biol. 126, 197–215 (2015).

10. L. B. Chen, Mitochondrial membrane potential in living cells. Annu. Rev. Cell Biol. 4, 155–181 (1988).

11. S. Meeusen, J. M. McCaffery, J. Nunnari, Mitochondrial fusion intermediates revealed in vitro. Science (80-.). 305, 1747–1752 (2004).

12. R. Ramzan, K. Staniek, B. Kadenbach, S. Vogt, Mitochondrial respiration and membrane potential are regulated by the allosteric ATP-inhibition of cytochrome c oxidase. Biochim. Biophys. Acta. 1797, 1672–1680 (2010).

13. P. K. Mouli, G. Twig, O. S. Shirihai, Frequency and selectivity of mitochondrial fusion are key to its quality maintenance function. Biophys. J. 96, 3509–3518 (2009).

14. W. Wang et al., A voltage-dependent K+ channel in the lysosome is required for refilling lysosomal Ca2+ stores. J. Cell Biol. 216, 1715–1730 (2017).

15. X. Wang et al., TPC proteins are phosphoinositide-activated sodium-selective ion channels in endosomes and lysosomes. Cell. 151, 372–383 (2012).

16. B. A. Schröder, C. Wrocklage, A. Hasilik, P. Saftig, The proteome of lysosomes. Proteomics. 10, 4053–4076 (2010).

17. H. Xu, E. Martinoia, I. Szabo, Organellar channels and transporters. Cell Calcium. 58, 1–10 (2015).

18. E. W. Miller et al., Optically monitoring voltage in neurons by photo-induced electron transfer through molecular wires. Proc. Natl. Acad. Sci. USA. 109, 2114–2119 (2012).

19. P. Yan et al., Palette of fluorinated voltage-sensitive hemicyanine dyes. Proc. Natl. Acad. Sci. USA. 109, 20443–20448 (2012).

20. E. W. Miller, Small molecule fluorescent voltage indicators for studying membrane potential. Curr. Opin. Chem. Biol. 33, 74–80 (2016).

21. J. Srivastava, D. L. Barber, M. P. Jacobson, Intracellular pH sensors: design principles and functional significance. Physiology (Bethesda). 22, 30–39 (2007).

22. H. H. Yang, F. St-Pierre, Genetically encoded voltage indicators: opportunities and challenges. J. Neurosci. 36, 9977–9989 (2016).

23. S. Surana, J. M. Bhat, S. P. Koushika, Y. Krishnan, An autonomous DNA nanomachine maps spatiotemporal pH changes in a multicellular living organism. Nat Commun. 2, 340 (2011).

24. S. Modi, C. Nizak, S. Surana, S. Halder, Y. Krishnan, Two DNA nanomachines map pH changes along intersecting endocytic pathways inside the same cell. Nat. Nanotechnol. 8, 459–467 (2013).

25. S. Modi et al., A DNA nanomachine that maps spatial and temporal pH changes inside living cells. Nat. Nanotechnol. 4, 325–330 (2009).

26. S. Saha, V. Prakash, S. Halder, K. Chakraborty, Y. Krishnan, A pH-independent DNA nanodevice for quantifying chloride transport in organelles of living cells. Nat. Nanotechnol. 10, 645–651 (2015).

27. K. Chakraborty, K. Leung, Y. Krishnan, High lumenal chloride in the lysosome is critical for lysosome function. Elife. 6, e28862 (2017).

28. B. Zhao et al., Visualizing Intercellular Tensile Forces by DNA-Based Membrane Molecular Probes. J. Am. Chem. Soc. 139, 18182–18185 (2017).

29. K. Chakraborty, A. T. Veetil, S. R. Jaffrey, Y. Krishnan, Nucleic Acid-Based Nanodevices in Biological Imaging. Annu. Rev. Biochem. 85, 349–373 (2016).

30. M. Famulok, J. S. Hartig, G. Mayer, Functional aptamers and aptazymes in biotechnology, diagnostics, and therapy. Chem. Rev. 107, 3715–3743 (2007).

31. S. Modi, S. Halder, C. Nizak, Y. Krishnan, Recombinant antibody mediated delivery of organelle-specific DNA pH sensors along endocytic pathways. Nanoscale. 6, 1144–1152 (2014).

32. D. Bhatia et al., Quantum dot-loaded monofunctionalized DNA icosahedra for single-particle tracking of endocytic pathways. Nat. Nanotechnol. 11, 1112–1119 (2016).

33. R. U. Kulkarni et al., Voltage-sensitive rhodol with enhanced two-photon brightness. Proc. Natl. Acad. Sci. USA. 114, 2813–2818 (2017).

34. J. Lahann, Ed., Click chemistry for biotechnology and materials science (John Wiley & Sons, Ltd, Chichester, UK, 2009).

35. H. Leonhardt, L. Gordon, R. Livingston, Acid-base equilibriums of fluorescein and 2’,7’-dichlorofluorescein in their ground and fluorescent states. J. Phys. Chem. 75, 245–249 (1971).

36. J. E. Whitaker et al., Fluorescent rhodol derivatives: versatile, photostable labels and tracers. Anal. Biochem. 207, 267–279 (1992).

37. B. van Lengerich, R. J. Rawle, S. G. Boxer, Covalent attachment of lipid vesicles to a fluid-supported bilayer allows observation of DNA-mediated vesicle interactions. Langmuir. 26, 8666–8672 (2010).

38. Y. Krishnan, M. Bathe, Designer nucleic acids to probe and program the cell. Trends Cell Biol. 22, 624–633 (2012).

39. M. J. Shattock, H. Matsuura, Measurement of Na(+)-K+ pump current in isolated rabbit ventricular myocytes using the whole-cell voltage-clamp technique. Inhibition of the pump by oxidant stress. Circ. Res. 72, 91–101 (1993).

40. R. Fliegert et al., Modulation of Ca2+ entry and plasma membrane potential by human TRPM4b. FEBS J. 274, 704–713 (2007).

41. S. R. Post et al., Class A scavenger receptors mediate cell adhesion via activation of G(i/o) and formation of focal adhesion complexes. J. Lipid Res. 43, 1829–1836 (2002).

42. A. Guha, V. Sriram, K. S. Krishnan, S. Mayor, Shibire mutations reveal distinct dynamin-independent and -dependent endocytic pathways in primary cultures of Drosophila hemocytes. J. Cell Sci. 116, 3373–3386 (2003).

43. J. L. Rosenfeld et al., Lysosome proteins are redistributed during expression of a GTP-hydrolysis-defective rab5a. J. Cell Sci. 114, 4499–4508 (2001).

44. J. G. Magadán, M. A. Barbieri, R. Mesa, P. D. Stahl, L. S. Mayorga, Rab22a regulates the sorting of transferrin to recycling endosomes. Mol. Cell. Biol. 26, 2595–2614 (2006).

45. M. P. Bradley, D. G. Rayns, I. T. Forrester, Effects of filipin, digitonin, and polymyxin B on plasma membrane of ram spermatozoa—an EM study. Arch Androl. 4, 195–204 (1980).

46. D. J. Yamashiro, S. R. Fluss, F. R. Maxfield, Acidification of endocytic vesicles by an ATP-dependent proton pump. J. Cell Biol. 97, 929–934 (1983).

47. J. Małecki, A. Wiedłocha, J. Wesche, S. Olsnes, Vesicle transmembrane potential is required for translocation to the cytosol of externally added FGF-1. EMBO J. 21, 4480–4490 (2002).

48. R. G. Johnson, A. Scarpa, Protonmotive force and catecholamine transport in isolated chromaffin granules. J. Biol. Chem. 254, 3750–3760 (1979).

49. M. Koivusalo, B. E. Steinberg, D. Mason, S. Grinstein, In situ measurement of the electrical potential across the lysosomal membrane using FRET. Traffic. 12, 972–982 (2011).

50. B. P. Crider, X. S. Xie, D. K. Stone, Bafilomycin inhibits proton flow through the H+ channel of vacuolar proton pumps. J. Biol. Chem. 269, 17379–17381 (1994).

51. Z. Farsi et al., Single-vesicle imaging reveals different transport mechanisms between glutamatergic and GABAergic vesicles. Science (80-.). 351, 981–984 (2016).

52. E. Lloyd-Evans, H. Waller-Evans, K. Peterneva, F. M. Platt, Endolysosomal calcium regulation and disease. Biochem. Soc. Trans. 38, 1458–1464 (2010).

53. S. Hover et al., Bunyavirus requirement for endosomal K+ reveals new roles of cellular ion channels during infection. PLoS Pathog. 14, e1006845 (2018).

54. C. C. Cain, D. M. Sipe, R. F. Murphy, Regulation of endocytic pH by the Na+,K+-ATPase in living cells. Proc. Natl. Acad. Sci. USA. 86, 544–548 (1989).

55. W. W. Douglas, T. Kanno, S. R. Sampson, Influence of the ionic environment on the membrane potential of adrenal chromaffin cells and on the depolarizing effect of acetylcholine. J. Physiol. (Lond). 191, 107–121 (1967).

56. R. Gagescu et al., The recycling endosome of Madin-Darby canine kidney cells is a mildly acidic compartment rich in raft components. Mol. Biol. Cell. 11, 2775–2791 (2000).

57. F. B. Schapiro, S. Grinstein, Determinants of the pH of the Golgi complex. J. Biol. Chem. 275, 21025–21032 (2000).

58. M. Liang et al., Identification of a pool of non-pumping Na/K-ATPase. J. Biol. Chem. 282, 10585–10593 (2007).

59. S. Gonin et al., Cyclic AMP increases cell surface expression of functional Na,K-ATPase units in mammalian cortical collecting duct principal cells. Mol. Biol. Cell. 12, 255–264 (2001).

60. M. Numata, J. Orlowski, Molecular cloning and characterization of a novel (Na+,K+)/H+ exchanger localized to the trans-Golgi network. J. Biol. Chem. 276, 17387–17394 (2001).

61. R. A. Saxton, D. M. Sabatini, mTOR Signaling in Growth, Metabolism, and Disease. Cell. 168, 960–976 (2017).

62. R. Puertollano, mTOR and lysosome regulation. F1000Prime Rep. 6, 52 (2014).

63. D. L. Medina et al., Lysosomal calcium signalling regulates autophagy through calcineurin and TFEB. Nat. Cell Biol. 17, 288–299 (2015).

64. X. Zhang et al., MCOLN1 is a ROS sensor in lysosomes that regulates autophagy. Nat Commun. 7, 12109 (2016).

65. C. Cang et al., mTOR regulates lysosomal ATP-sensitive two-pore Na(+) channels to adapt to metabolic state. Cell. 152, 778–790 (2013).

66. C. Cang, B. Bekele, D. Ren, The voltage-gated sodium channel TPC1 confers endolysosomal excitability. Nat. Chem. Biol. 10, 463–469 (2014).

67. P.-M. Wong, Y. Feng, J. Wang, R. Shi, X. Jiang, Regulation of autophagy by coordinated action of mTORC1 and protein phosphatase 2A. Nat Commun. 6, 8048 (2015).

68. L. N. Hockey et al., Dysregulation of lysosomal morphology by pathogenic LRRK2 is corrected by TPC2 inhibition. J. Cell Sci. 128, 232–238 (2015).

69. Q. Cao et al., BK channels alleviate lysosomal storage diseases by providing positive feedback regulation of lysosomal ca2+ release. Dev. Cell. 33, 427–441 (2015).

70. U. S. Lee, J. Cui, BK channel activation: structural and functional insights. Trends Neurosci. 33, 415–423 (2010).

71. S. P. Olesen, E. Munch, P. Moldt, J. Drejer, Selective activation of Ca(2+)-dependent K+ channels by novel benzimidazolone. Eur. J. Pharmacol. 251, 53–59 (1994).

72. D. Shen et al., Lipid storage disorders block lysosomal trafficking by inhibiting a TRP channel and lysosomal calcium release. Nat Commun. 3, 731 (2012).

73. M. Abu-Remaileh et al., Lysosomal metabolomics reveals V-ATPase- and mTOR-dependent regulation of amino acid efflux from lysosomes. Science (80-.). 358, 807–813 (2017).

74. G. A. Wyant et al., mTORC1 Activator SLC38A9 Is Required to Efflux Essential Amino Acids from Lysosomes and Use Protein as a Nutrient. Cell. 171, 642–654.e12 (2017).

