## Supplementary information for "A DNA-based voltmeter for organelles"

**This PDF file includes:**

Materials and Methods
Supplementary text
Figs. S1 to S21
Tables S1 to S2
Captions for Movies S1 to S2
References

**Other Supplementary Materials for this manuscript include the following:**

Movies S1 to S2

### Materials and Methods

Reagents

All modified oligonucleotides (Table S1) were purchased from IDT (USA). Fluorescently labelled oligonucleotides were subjected to ethanol precipitation and quantified by UV absorbance at 260nm. CellMask$^{TM}$ reagents and TMR-Dextran were purchased from molecular probes/Life Technologies (USA). Maleylated BSA (mBSA) and fluorescent transferrin (Tf-Alexa546) were conjugated according to previously published protocols (*1*, *2*). Torin-1, ML-SA1, NS1619 and *trans*-Ned-19 were purchased from Cayman Chemical (USA). Phenyl triflimide was purchased from TCI America (USA). N-Boc-piperazine was purchased from Oakwood chemicals (USA). Azido-PEG4-NHS was purchased from Click Chemistry tools (USA). 1-palmitoyl-2-oleoyl-sn-glycero-3-phosphoethanolamine lipid was purchased from Avanti lipids (USA). All other reagents were purchased from Sigma-Aldrich (USA) unless otherwise specified.

Synthesis of RVF

Synthesis scheme shown in Fig. S1. Compound 1a (0.2 g, 0.36 mmol) was suspended in N,N-dimethylformamide (DMF) (1.4 mL) and cooled to 0°C. Triethylamine (2.7 mL) and Phenyl triflimide (0.25 g, 0.72 mmol, 2 eq.) were added dropwise. The ice bath was removed and the reaction was stirred at room temperature for 2 hours. The reaction mixture was then diluted in water and extracted with dichloromethane (DCM) twice. Organic extracts were washed three times with brine and 1M HCl. The product was concentrated via rotor evaporation and dry DMSO was added (2mL). N-boc piperazine (3.72g, 20 mmol, 100eq.) was added and the reaction mixture was kept at 100ºC overnight. The reaction mixture was then diluted in water, extracted in DCM twice then washed with brine twice. The mixture was dried over anhydrous $Na_2SO_4$ and concentrated under reduced pressure. Silica gel column chromatography was performed in 5% methanol in chloroform to get 1b (0.08g, 28%). ESI-MS (-) Expected mass = 728.98, found = 729.0.

A reaction tube was charged with 1b (70 mg, 0.0956 mmol), $Pd(OAc)_2$ (6.87 mg, 0.0306 mmol, 0.32 eq.), tri-o-tolylphosphine (20.4 mg, 0.067 mmol, 0.7 eq.) and (*E*)-N,N-dimethyl-4-(4-vinylstyryl)aniline (26.23mg, 1.052 mmol, 1.1 eq.) which was previously synthesized according to literature (*3*). The tube was evacuated and backfilled with $N_2$ three times. 1 mL of dry DMF and 500 µL dry triethylamine were added via syringe, and the reaction was stirred at 110ºC overnight. The reaction mixture was diluted in water and extracted with DCM twice. The organic extract was washed with brine twice and concentrated under reduce pressure. Silica gel column chromatography was performed in 5% methanol in DCM to get orangish brown solid 1c (20 mg, 25%). ESI-MS (+), Expected mass = 851.29, found = 851.2.

A reaction vial with 1c (5 mg, 6.7 nmol) was placed in 5% TFA in DCM overnight for deprotection. TFA was removed under reduced pressure and Azido-PEG4-NHS ester (26mg, 6.7 nmol, 10.0 eq.), 300 µL dry DMF, and 200 µL triethylamine were added. The mixture was stirred for 4 hours at room temperature. The reaction mixture was then diluted in water, extracted to DCM twice, washed with brine three times, dried over anhydrous $Na_2SO_4$ and concentrated under reduced pressure. Silica gel column chromatography was performed with 2% methanol in DCM, slowly increasing the gradient to 10%, to get reddish brown solid 1d (2mg, 30%). ESI-MS(+), Expected mass = 1025.23, found = 1025.3.

Sample preparation

See table S1 for detailed sequence information.

Sensing domain ($D_V$, $D_V^{RE}$): DBCO modifed single stranded DNA was conjugated to RVF using a previously reported copper free click chemistry protocol (Fig. S2) (*4*). 10 µM of the 3'- DBCO modified 38 base strand (IDT, USA) was coupled to the azide containing RVF (50 µM, 5 eq.) in 20 mM sodium phosphate buffer, pH 7.4. The reaction mixture was stirred for 4 hours at room temperature (RT). Upon completion, unconjugated fluorophores were removed by ethanol precipitation (*5*). RVF conjugated DNA was reconstituted in 10 mM phosphate buffer, pH 7.4 and the final concentration was quantified using UV absorbance at 260 nm.

Targeting domain ($D_T$): 1-palmitoyl-2-oleoyl-sn-glycero-3-phosphoethanolamine (POPE) was conjugated to NHS-PEG4-Azide using an established protocol (*6*). 20 µM of the 5'-DBCO modified 22 base strand (IDT, USA) was coupled to azido-POPE (40 µM, 2eq.) in 20 mM sodium phosphate buffer, pH 7.4 and stirred for 4 hours at RT (Fig. S2).

Construction of *Voltair*$^{PM}$, *Voltair*$^{IM}$ and *Voltair*$^{RE}$: Stock solution of *Voltair*$^{PM}$ was prepared at a final concentration of 10 µM by mixing $D_V$, $D_T$ and $D_A$ (Atto647N – 5' modified strand) at an equimolar ratio in 10 mM sodium phosphate buffer, pH 7.4 (Fig. S2, S3). For *Voltair*$^{IM}$ samples, $D_V$ and $D_{A'}$ (Atto647N – 3' modified strand) were mixed at an equimolar ratio with a final concentration of 10 µM. For *Voltair*$^{RE}$ samples, 10 µM of $D_V^{RE}$, $D_{Tf}$ and $D_A^{RE}$ were mixed at an equimolar ratio. For all samples, annealing was performed by heating the reaction mixture to 95°C for 15 min and gradually cooling to RT, at 1°C/ 3 min (*7*). Annealed samples were equilibrated at 4°C overnight.

Gel electrophoresis:

For gel electrophoresis 15% native polyacrylamide gels were used for annealed samples. Denaturing polyacrylamide gels containing 12-15% acrylamide [38:2 acrylamide/ bisacrylamide] were used for dye conjugated single strand samples. Gels were run in 1X TBE buffer (100 mM Tris. HCl, 90 mM boric acid and 2 mM EDTA, pH 8.3) at RT. Non-fluorescent samples were stained with ethidium bromide (1µg/mL) for 10 mins prior to visualization. Samples were observed by Biorad Universal Hood II Gel Doc system (Bio-Rad Laboratories, Inc.)

In vitro spectroscopic measurements:

Fluorescence spectra were recorded on a FluoroMax-4 scanning Spectro-fluorometer (Horiba Scientific, Edison, NJ, USA). The *Voltair*$^{IM}$ was diluted to 100 nM in universal buffer, UB4 buffer (20 mM HEPES, MES and sodium acetate, 150 mM KCl, 5 mM NaCl, 1 mM $CaCl_2$ and $MgCl_2$) of desired pH for all in vitro fluorescence experiments. For recording spectra, *Voltair*$^{IM}$ samples were excited at 520 nm and 650 nm and emission spectra were collected between 525 – 600 nm and 655-750 nm respectively. In order to study the pH sensitivity of *Voltair* probes, buffers of indicated pH (4 – 7) were incubated with *Voltair*$^{IM}$ for 30 mins prior recording. The pH sensitivity was obtained by plotting the ratio of RVF emission intensity (G, $\lambda_{Ex}$ = 520 nm) at 550 nm and emission intensity of normalizing dye (R, $\lambda_{Ex}$ = 650 nm) at 665 nm, as a function of pH (Fig. S5). Mean of G/R ratio from three independent experiments and their standard deviation were plotted for each pH value.

Cell culture, plasmids and transfection:

Human embryonic kidney cells (HEK 293T) were a kind gift from Prof. Bryan Dickinson's lab at the University of Chicago. Cells were cultured in Dulbecco's Modified Eagle's Medium (Invitrogen Corporation, USA) containing 10% heat inactivated Fetal Bovine Serum (FBS) (Invitrogen Corporation, USA), 100 U/mL penicillin and 100 µg/mL streptomycin and maintained at 37ºC under 5% $CO_2$. HEK 293T cells were passaged and plated at a confluency of 20 – 30% for electrophysiology experiments, and 50 – 70% for transfection and intracellular measurements.

The hMSR1 sequence was cloned into the PCS2NXE vector (4,103 bp) containing the CMV promoter for overexpression in mammalian cell lines. The hMSR1-CFP plasmid (5973 bp) was constructed by cloning hMSR1 sequence into pECFP-C1 plasmid (4731 bp). The identity of each construct was confirmed by sequencing, using forward primer (5' to 3') GGGACATGGGAATGCAATAG and reverse primer (5' to 3') CTCAAGGTCTGAGAATGTTCCC.

The mCherry-TGNP-N-10 was a gift from Michael Davidson (Addgene plasmid #55145) and Rab7-RFP was a gift from Ari Helenius (Addgene plasmid #14436, (*8*)). Construction of ScFv-Furin construct is reported previously (*9*).

HEK 293T cells were transiently transfected with respective plasmids using *Trans*IT®-293 transfection reagent (MIRUS). After a 4-hour incubation the transfected medium was replaced with fresh medium. Labeling experiments were performed on cells 48 hours post transfection.

Electrophysiology:

A schematic of the electrophysiology equipment used for whole cell patch clamp recording is shown in Fig. S7. Recordings of probe labelled HEK 293T cells were performed with an Axopatch 200A amplifier (Molecular Devices). The signals were digitized using an NI-6251 DAQ (National Instruments). The amplifier and digitizer were controlled using WinWCP software (Strathclyde Electrophysiology Software).

Patch pipettes were pulled using a Sutter P-97 Micropipette puller. Borosilicate glass capillaries (Sutter) of dimension 1.5 mm x 0.86 mm (OD/ID) were pulled using the program: Heat – Ramp, Pull – 0, Vel – 21, Time – 1(Delay), Loops – 5. Patch pipettes with resistances between 5-10 MOhm were used in voltage clamping experiments. The patch pipette was positioned using an MP325 motorized manipulator (Sutter). Image Acquisition software Metamorph premier Ver. 7.8.12.0 was linked to an NI-6501 DAQ to enable voltage triggered image acquisition. When applying a voltage pulse across the cell membrane a digital output pulse was generated by WinWCP to trigger imaging. For all measurements the extracellular solution composition was 145 mM NaCl, 20 mM glucose, 10 mM HEPES, pH 7.4, 3 mM KCl, 2 mM $CaCl_2$, 1 mM $MgCl_2$ (310 mOsm) and the intracellular solution composition was 115 mM potassium gluconate, 10 mM EGTA tetrapotassium salt, 10 mM HEPES, pH 7.2, 5 mM NaCl, 10 mM KCl, 2 mM ATP disodium salt, 300 µM GTP trisodium salt (290 mOsm).

For plasma membrane voltage clamping experiments, 1 µM RVF or 500 nM *Voltair$^{PM}$* was incubated with HEK 293T cells for 30 mins in Hank's Balanced Salt Solution (Thermofisher) for plasma membrane labeling at room temperature. Labelled cells were washed three times with 1X PBS and incubated in extracellular solution for whole cell voltage clamping. Whole cell voltage clamping was performed according to an established protocol (*10*). Electrophysiological measurements were made from a single HEK 293T cell, without physical interaction with nearby cells to avoid interference from gap junctions. For background subtraction, bleaching correction and lamp fluctuation compensation, imaging field was chosen with at least one more HEK 293T cell that is not clamped. Once clamped, membrane potential is changed from -100 mV to +100 mV in 10mV increments at 1000 ms intervals. Around 200 ms after the voltage is changed three images are taken in quick succession. Voltage clamp experiments were also performed with extracellular solutions of different pH, to study the effect of pH on voltage sensitivity of RVF (Fig. S5).

Microscopy:

Wide field microscopy was carried out on an IX83 inverted microscope (Olympus Corporation of the Americas, Center Valley, PA, USA) using either a 100X or 60X, 1.4 NA, DIC oil immersion objective (PLAPON, Olympus) and Evolve Delta 512 EMCCD camera (Photometrics, USA). Filter wheel, shutter and CCD camera were controlled using Metamorph premier Ver 7.8.12.0 (Molecular Devices, LLC, USA). Images on the same day were acquired under the same acquisition settings (exposure 100 ms and EM gain at 100 for Atto647N, exposure 200 ms and EM gain 300 for RVF). All the images were background subtracted by taking mean intensity over an adjacent cell free area. The mean intensity in each endosome/lysosome was measured in the sensing channel (G) and the normalizing channel (R). Filter sets, purchased from Chroma, suitable for each fluorophore were selected to minimize excitation and emission from other dyes in the sample. RVF channel images were obtained using 500/20 band pass excitation filter, 535/30 band pas emission filter and 89016 dichroic. For Atto647N, images were obtained using the 640/30 band pass excitation filter, 705/72 band pass emission filter and 89016 dichroic. Pseudo-color images were generated by calculating the G/R ratio per pixel in Fiji using the Image calculator module. For ratiometric pH measurements, fluorescence of FITC-dextran was recorded by exciting at 480 nm and 430 nm and collecting emission at 520 nm for both images. Images were acquired using a 480/20 or 430/24 band pass excitation filter, 520/40 band pass emission filter and 89016 dichroic.

Intracellular membrane potential measurements were performed by recording images in the sensing channel (G) and the normalizing channel (R) in cells where each specific compartment was labeled. After acquisition of G and R images, intracellular membrane potential was neutralized by adding 50 μM valinomycin and monensin for 20 mins at room temperature. A set of G and R images of same cells were acquired after valinomycin and monensin or pharmacological treatments as shown in Fig. S16. These images of neutralized endosomes were used to as a baseline measurement to correct for variations in autofluorescence. To record all endo-lysosomal compartments in the cell, Z-stacks (30 planes, Z distance = 0.8 μm) were captured and a maximum intensity projection was used to produce a single image for analysis. Confocal images were captured with a Leica TCS SP5 II STED laser scanning confocal microscope (Leica Microsystems, Inc. Buffalo Grove, IL, USA) equipped with a 63X, 1.4 NA, Oil immersion objective. RVF was excited using an argon laser with 514 nm wavelength, CFP by 458 nm and Atto647N using a He-Ne laser with 633 nm wavelength. CellMask orange stain was excited by 543 nm and all emissions were filtered using Acousto Optical Beam Splitter (AOBS) with settings suitable for each fluorophore and recorded using hybrid detectors (HyD).

Competition assay:

HEK 293T cells transfected with hMSR1-CFP were washed with 1X PBS, pH 7.4 prior to labeling with the DNA device. Cells were incubated with 20 μM maleylated BSA (mBSA) for 15 min, followed by a 20 min pulse of 500 nM DNA device and 20 μM mBSA to allow internalization by receptor mediated endocytosis. Control cells were washed and pulsed for 20 mins in 500 nM DNA device without mBSA. Cells were then washed three times with 1X PBS and chased for 30 mins in complete DMEM media. Complete media was replaced with Opti-MEM solution (Thermofisher) and imaged by widefield microscopy for quantification and confocal microscopy for representative images. Whole cell intensities in the Atto647N channel were quantified for cells expressing hMSR1-CFP observed in CFP channel. The mean intensity from three different experiments were normalized against the no mBSA control, for n~50 cells.

Co-localization and labeling experiments:
In order to find out time points when internalized DNA devices specifically label different intracellular organelles, we performed colocalization experiments with different organelle specific markers as a function of time in HEK 293T cells. Transferrin receptors are known to recycle to plasma membrane via early endosome (EE) and recycling endosome (RE), therefore we used fluorescent Transferrin to specifically label EE by pulsing it for 10 minute prior to imaging and label RE with an additional chase time of 30 mins (Fig S20) (*11*). Rab7-RFP is a well-established late endosome (LE) marker, and was transiently expressed in HEK 293T cells to label LE (*12*). Transient expression of TGN46-mCherry specifically labels trans Golgi network (TGN) (*13*). Finally, TMR-Dextran a specific marker for lysosome (Ly) after previously reported chase times, was pulsed for 1 hour, followed by 16 hours chase in complete media to label lysosomes (*12*).
To find out the trafficking time of DNA device in specifically labeling EE, HEK 293T cells transfected with hMSR1, were pulsed with 500 nM DNA device containing Atto647N for 10 minutes and chased for indicated time. These cells were also pulsed with 100 nM Tf-Alexa546 for 10 mins to visualize the early endosomes. A stock solution of Tf-Alexa546 was prepared according to previously established methods (*1*). Images of the Alexa546 channel and the Atto647N channel were acquired by confocal imaging (microscopy methods) and Pearson correlation coefficient (PCC) value were calculated using Fiji plugin coloc2 (*14*).
To find out the trafficking time of DNA device in specifically labeling LE or Ly, HEK 293T-hMSR1 cells were transfected with Rab7-RFP for LE or pre-pulsed & chased with TMR-Dextran for Ly. These cells were then pulsed with 500 nM DNA device for 30 mins and chased for indicated time. PCC of DNA device colocalization with EE, LE or Ly was plotted with respect to chase time. A maximum PCC value represents the time point at which DNA device is localized to the specific organelle. The trafficking time of DNA device with transferrin aptamer and $d(AT)_4$ tag in labeling RE and TGN, respectively, have been established previously (*1*, *9*). Briefly, recycling endosomes are targeted by pulsing *Voltair*$^{RE}$ in 1X HBSS, for 10 mins at 37°C, followed by 30 mins of chase in complete media at 37°C. Trans Golgi network is targeted by pulsing *Voltair*$^{IM}$ in complete media containing cycloheximide (CHX) for 90 mins at 37°C, followed by 90 mins chase in the same media.

Intracellular voltage clamping:
Ionophores selectively permeabilize specific ions across the membrane and have been used to manipulate the membrane potential of purified lysosomes (*15*). Three well established ionophore sets have been used previously, (i) Valinomycin, a $K^+$ ionophore which gives the lysosome a membrane potential of +40 mV when incubated for 10 mins at RT, (ii) An equimolar concentration of valinomycin and monensin, in presence of 150 mM $K^+$ which neutralizes lysosomal membrane potential. (iii) FCCP, a $H^+$ ionophore which in absence of ATP shifts the lysosomal membrane potential to -90 mV (*15*). Lysosomes of HEK 293T cells were labelled

with 500 nM *Voltair*$^{IM}$ as previously described above. Cells were treated with 10 μM digitonin for 2 mins in presence of an intracellular solution with one set of ionophores and washed three times with the same buffer in absence of digitonin. This replaces the cytosol with the intracellular solution containing ionophores which clamp lysosome voltage to known values. Pseudo color images shown in fig S15, were generated as described in image analysis section. Ratiometric G and R images were acquired to record G/R values of ~50 lysosomes for each experiment and plotted as a function of membrane potential (Fig 3C).

Image analysis:
Images were analyzed with Fiji or imageJ (NIH, USA). For organellar voltage measurements, regions of cells containing single isolated endosomes/lysosomes in each Atto647N (R) image were manually selected and the coordinates saved in the ROI plugin in ImageJ. Similarly, for background computation, a nearby region outside endosomes/lysosomes were manually selected and saved as an ROI. The same regions were selected in the RVF (G) image by recalling the ROIs. After background subtraction, mean intensity for each endosome (G and R) was measured and exported to OriginPro (OriginLab, USA). A ratio of G to R intensities (G/R) was obtained from these values by dividing the mean intensity of a given endosome in the G image with the corresponding intensity in the R image. We used the same protocol for individual endosomes in the calibration, therefore G/R ratio correspond to membrane potential calibrated intracellularly. To minimize the measurement error due to low fold change of *Voltair* probes, the same endosomes or lysosomes are measured post addition of valinomycin and monensin which neutralizes the membrane potential in presence of 150 mM KCl. For TGN voltage measurements, total cell intensity was recorded and background subtraction was performed by manually selecting a region outside the cell. For a given experiment, membrane potential of an organelle population was determined by converting the mean $[G/R]_V$ - $[G/R]_O$ value of the distribution to voltage values according to intracellular voltage calibration profile (Fig. S17). The mean value of each organelle population across three trials on different days is determined and the final data is presented as mean ± S.E.M. Representative images are shown in pseudo-color images, where G and R images were modified by thresholding in ImageJ to get G' and R' images. Using ImageJ's Image calculator module, G' images were divided by R' images to generate an image where each pixel represents $[G/R]_V$.
For pH measurements in lysosomes of digitonin treated cells, images acquired by 430 excitation and 480 excitations were background subtracted and individual lysosomes were selected as mentioned above. The average ratio of 480 to 430 excitation for the population of lysosomes was computed in Origin. Representative pseudo-color images were constructed as explained above. For whole cell patch clamp, image analysis was performed using custom Matlab code. A series of images corresponding to a voltage sweep from -100 to +100 mV was collected and input into the program. By identifying changes in intensity from the first to last image a region of interest corresponding to the clamped cell was selected. Other cells present in the image were also selected based on an intensity threshold. The region containing no cells was used to subtract background noise from the detector from all regions. Intensity from unclamped cells was measured in each image and used to correct for photobleaching or fluctuation in lamp intensity. After background corrections the average intensity of the patch clamped cell was then measured for each individual image in the series and normalized against the value at -60 mV. This analysis code will be shared upon request.

Statistical analysis

For statistical analysis between two samples, two-sample two tailed test assuming unequal variance were used. For comparison of multiple samples, one-way ANOVA with a post hoc Tukey test or Fischer test was used. All statistical analysis was performed in Origin (Student version). In box plots, the mean, median line, 25-75% box and Standard deviation were provided.

**Supplementary text:**

Characterization of *Voltair* using gel electrophoresis.

Copper free click reaction of RVF to DBCO labeled strand ($D_V$) was validated by 15% Denaturing PAGE run in 1X TBE, at 150 V. Conjugation of 1 KDa (RVF) to 10 KDa (DBCO-strand) causes the slow mobility shift of $D_V$ strand in Fig. S3. Furthermore, we confirmed that the lower mobility band contains RVF by fluorescence imaging in the rhodamine channel (excited by Epi-light and filtered by 560DF50). The $D_V$ strand was purified and hybridized with the normalizing ($D_A$) and targeting module ($D_T$) as described in sample preparation section. A 15% native PAGE was run to characterize the formation of complete sensor. In 15% acrylamide gels there is a large shift between duplex DNA and ssDNA. Under these conditions the increased persistence length of dsDNA leads to much slower mobility of *Voltair*$^{PM}$ with respect to single strand components ($D_V$, $D_T$, $D_A$). This therefor validates the assembly, at very high yield (> 99%) of the full sensor. We again confirmed the slower mobility band contains RVF and the normalizing dye by imaging fluorescence in rhodamine channel (RVF) and Atto647N channel (Fig. S3).

pH insensitivity of *Voltair* probes

Endocytic compartments show increasing levels of lumenal acidity along the maturation pathway, ranging from pH 6.5 at EE to pH 4.5 at Ly in mammalian cells. Most fluorescein-based probes are sensitive to acidic pH. Protonation of phenolic -OH moiety (pKa = 6.4) of fluorescein-based chromophore decreases the fluorescence and hence used as well-established pH sensors. This aspect of pH interference makes it difficult to uncouple the probes ability to sense signals. Thus, we chose a voltage sensitive dye (RVF) that could reliably sense at acidic pH. RVF comprise of dichlorosulforhodol dye, which is expected to have very low pKa due to two reasons. (i) dichlorofluorescein derivatives have lower pKa (~4.5) than fluorescein due to the negative inductive effect of chloro substitution at the ortho position of the phenolic OH. (*16*). (ii) rhodol fluorophores have lower pKa than fluorescein (pKa of Rhodal = 5.5) (*17*). To confirm the pH independence of RVF in sensing voltage, HEK 293T cells were labeled with 1 μM RVF and voltage clamped from -100 mV to +100 mV with extracellular solutions of pH 4.5, 5 and 7 (Fig S5). To ensure pH insensitivity of complete sensor *Voltair*$^{IM}$, fluorescence spectra of RVF and Atto647N were recorded at different pH (4.0 – 7.0). G/R ratios were calculated by dividing the emission spectra maxima of RVF (550 nm) to that of Atto647N (665 nm) and plotted with respect to pH as shown in Fig. S5.

DNA-lipid conjugate labels plasma membrane:

To anchor DNA based probes to the outer membrane leaflet of cell, the DNA backbone was conjugated to 1-palmitoyl-2-oleoyl-sn-glycero-3-phosphoethanolamine lipid (POPE). POPE-DNA conjugates have been previously shown to effectively label the plasma membrane of cells

(*18*). For efficient insertion, the DNA duplex was coupled to POPE via tetraethylene glycol linker, providing additional flexibility and spacing (Fig S2, S6). DNA-lipid conjugates show reversibility in anchoring due to negatively charged DNA as a hydrophilic head group. We labeled the cells with 500 nM DNA-POPE or *Voltair$^{PM}$* for 20 mins at room temperature, washed the cells three times with 1X PBS and incubate the cells in extracellular buffer.

Digitonin mediated plasma membrane permeabilization assay:
To map the response characteristics of *Voltair$^{IM}$* in intracellular membranes, we developed a method to exchange the cytosol with a chosen buffer without disrupting the lysosomal membrane containing *Voltair$^{IM}$*. Digitonin is commonly used to permeabilize the cell membrane for cell biology techniques such as immunostaining and cell lysis. Transient application of digitonin has shown to selectively permeabilize the plasma membrane (PM) of cells, without affecting intracellular compartments (Fig. S13, S14) (*19*, *20*). To standardize and validate the technique we labeled lysosomes of HEK 293T cells with TMR-Dextran (10 KDa and ~2 nm diameter) and selectively permeabilized the PM with 10 μM digitonin in a desired buffer. The treatment creates pores with 30 nm diameter on the PM (*20*). Treatment with the lipophilic CellMask™ before and after digitonin treatment showed that Cellmask™ could access and label internal membranes only post-digitonin treatment, confirming the formation of pores in the PM. However, TMR dextran showed no leakage from the lysosomes, indicating that the lysosomal membrane had not been breached (Fig S13). Washing away the digitonin with the desired buffer reseals the PM, and thereby the cytosolic milieu is replaced with that of the buffer. To demonstrate this, lysosomes were labelled with pH-sensitive probe FITC Dextran. Dual excitation pH imaging in cultured cells revealed a low 480/430 ratio characteristics of acidic lysosomes. Upon digitonin treatment we replaced the cytosol of these cells with 20 mM HEPES buffer, pH 7.4 of intracellular ionic composition (IC buffer) lacking ATP. Without available ATP in the milieu, the ATP-dependent lysosomal proton pump V-ATPase, cannot pump in protons to acidify the lysosome lumen. As a result, the lysosomal pH increases as revealed by the high 480/430 ratio of FITC Dextran. However, a similar procedure using an IC buffer containing 2 mM ATP yields lysosomes of high acidity indicating that the integrity of the lysosome is preserved, the pumps and transporters are functional and that the membrane potential is actively maintained. ATP dependent organelle acidification is sensitive to $NH_4Cl$ which effectively neutralizes acidic compartments like the lysosomes. Accordingly, digitonin treatment with IC buffer containing 10 mM $NH_4Cl$ effectively neutralized lysosomes, indicated that this was a robust method to replace the cytosol with a buffer of desired composition without affecting organelle integrity.

Organellar membrane potential measurements:
Intracellular measurements of membrane potential are performed by calculating G/R values of single endo-lysosomes. The intracellular calibration plot yields the sensitivity slope as m = 0.00235 ± 2.96E-4, using which the following equation was formulated to calculate membrane potential.

$$V = K \times (1 - G/R); \quad K = 1/m = 425.5 \ \pm 53.5 \, mV;$$

The error in measurements is caused by two factors, (1) fluorescence measurement error in a recording G/R values (G/R ± ΔG/R) and (2) Calibration error due to the digitonin mediated calibration plot. Calibration error is calculated by considering the error in measurement of K. (K ± ΔK). Error analysis was performed to calculate the membrane potential error shown in Table S2.

*Voltair$^{IM}$* stability assay**:**

Endo-lysosomal nucleases have been reported to degraded nucleic acids trafficked to lysosomes (*21*, *22*). In order to reliably measure membrane potential, *Voltair$^{IM}$* probes must be stable at the time point when the measurement is made. We assessed the stability of *Voltair$^{IM}$* by recording the fluorescence signal of RVF inside lysosomes with respect to Atto647N signal. Degradation of *Voltair$^{IM}$* results in free RVF which docks in the inner leaflet and Atto647N which leaks out of the lysosome membrane (*23*). Therefore, the ratio of intensities should decrease as the DNA scaffold is degraded. Plotting G/R ratio of lysosomes with respect to time revealed that *Voltair$^{IM}$* probes are fully stable up to 140 min.

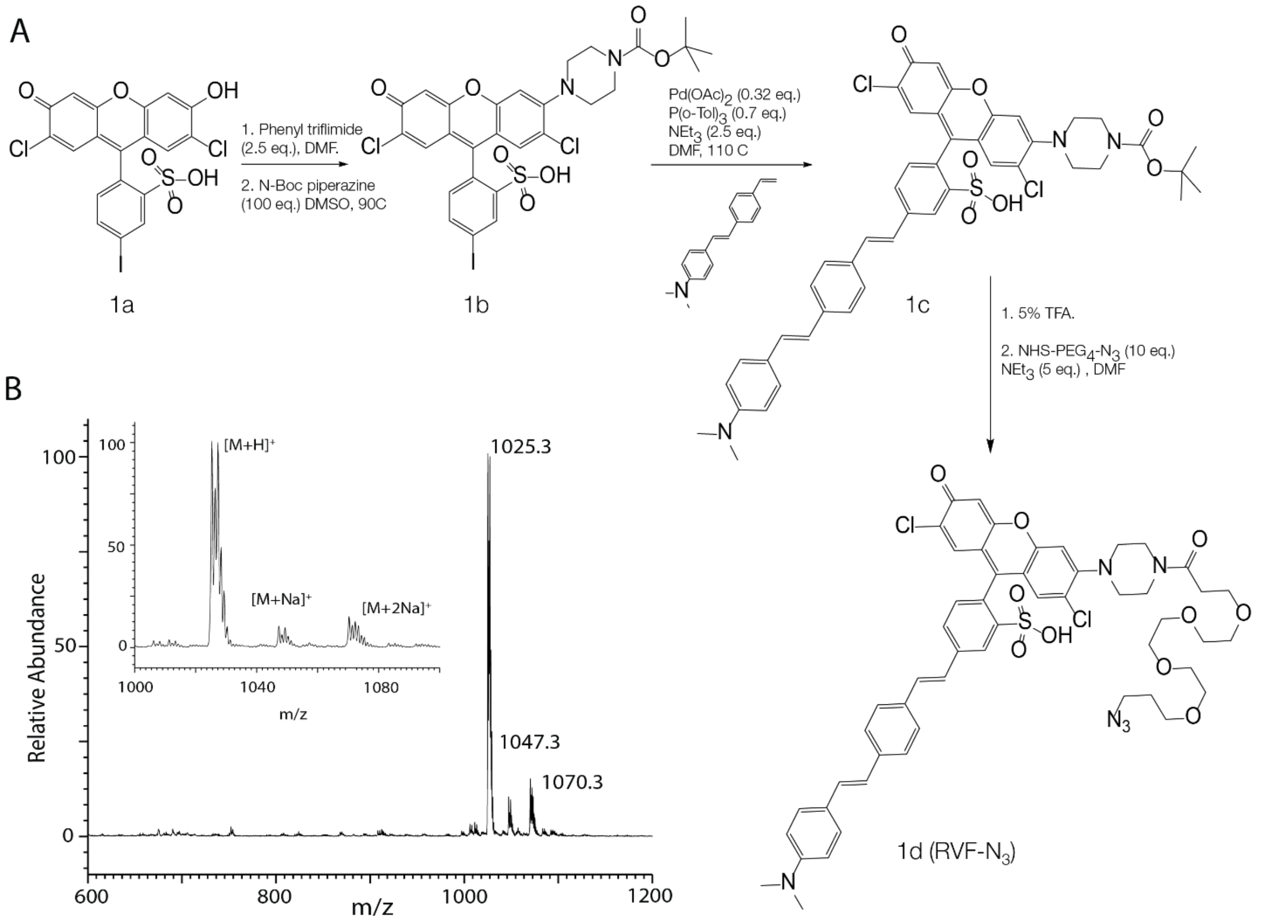


**Fig. S1. Synthesis of conjugatable RVF-$N_3$**. (A) Reaction scheme for synthesis of azido-RVF. The azido group in RVF is utilized to conjugate it to the DNA strand Dv, the sensing component in the *Voltair* probes. (B) ESI-MS of RVF-$N_3$. Inset shows the region corresponding to the M+2 peaks arising from the Cl isotopes.

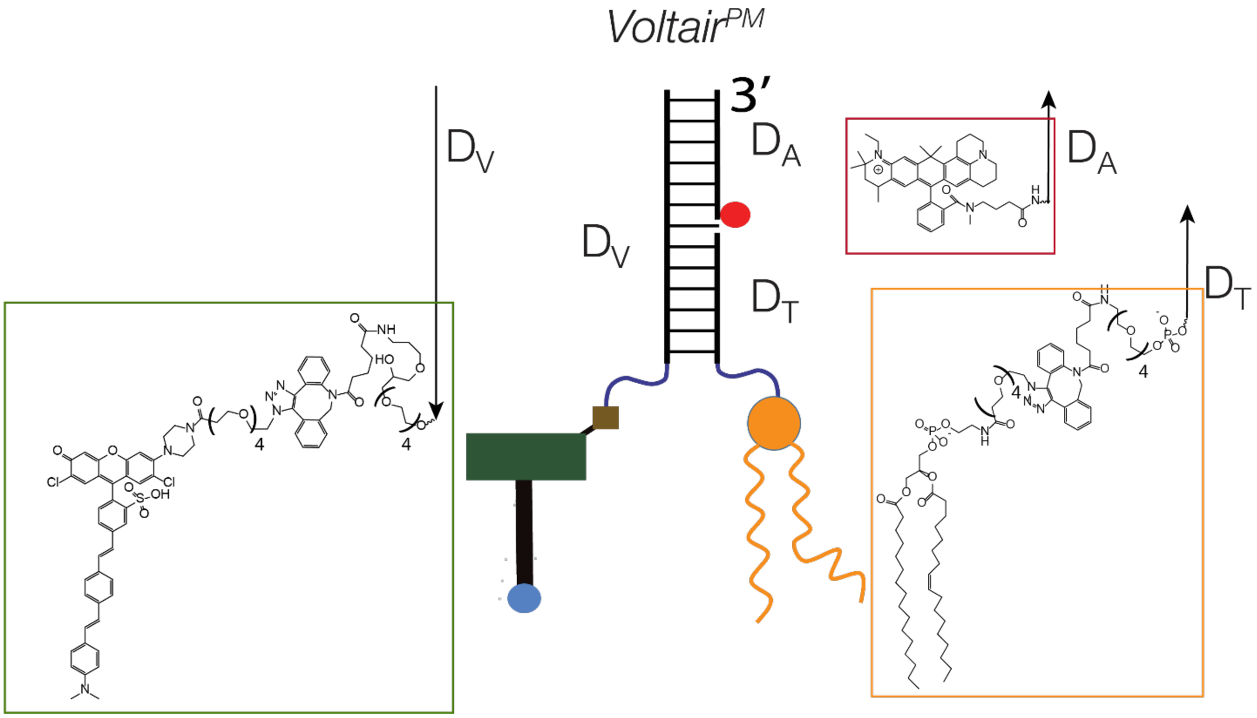


**Fig. S2. Components of *Voltair$^{PM}$*.** *Voltair$^{PM}$* is a trimeric complex comprising of voltage sensing strand $D_V$, normalizing strand $D_A$ and targeting strand $D_T$. $D_V$ is constructed by coupling RVF-$N_3$ to DBCO-ispacer18-DNA. $D_A$ strand is an Atto647N modified at 5' end of DNA. $D_T$ is constructed by coupling POPE-$N_3$ to DBCO labeled DNA. These individual strands are assembled to form the complete sensor.

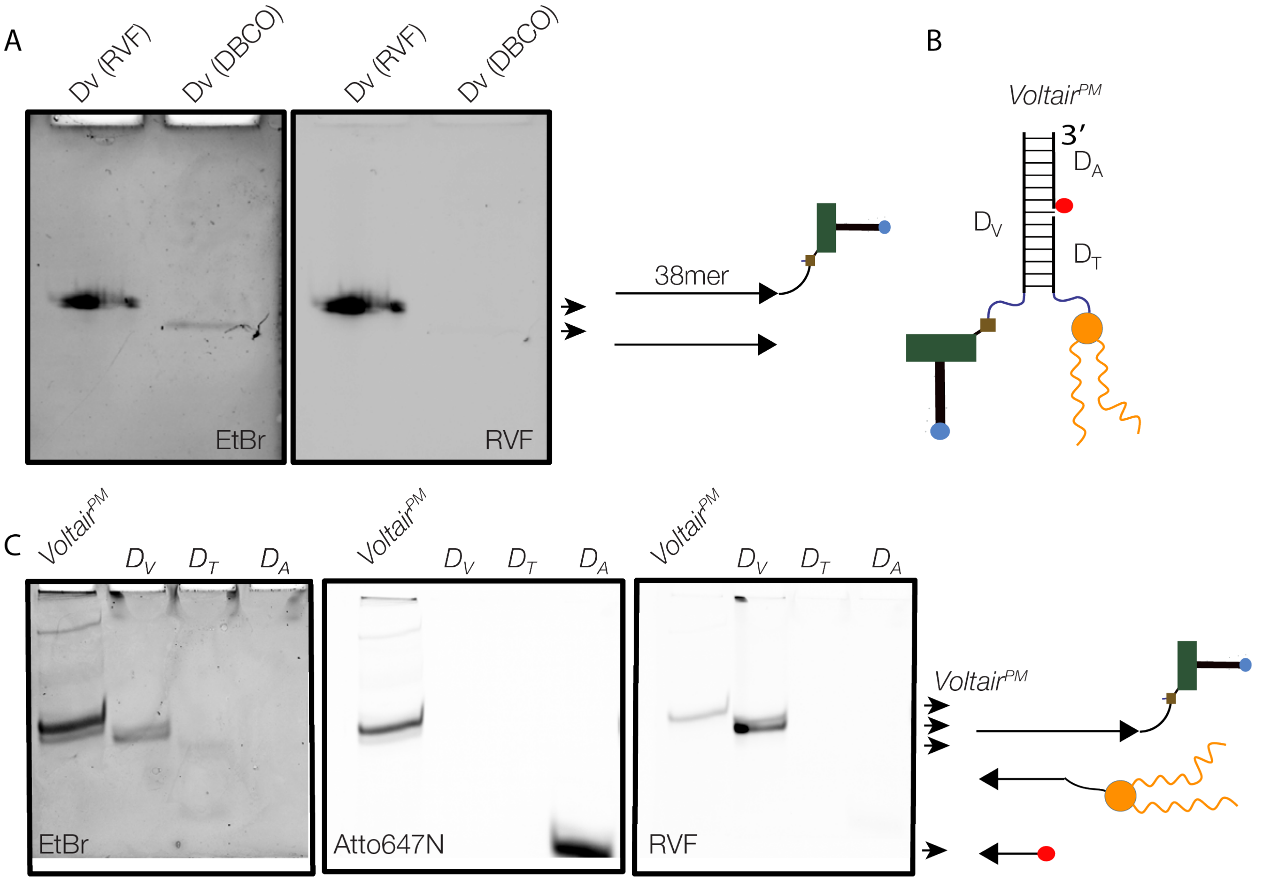


**Fig. S3. Gel characterization of *Voltair$^{PM}$*.** A) 15% Denaturing polyacrylamide gel electrophoresis in 1X TBE, showing the conjugation of RVF-$N_3$ to the DNA strand $D_V$ bearing a 3’ DBCO functionality. Gels are imaged in the EtBr channel that stains DNA as well as the rhodamine channel for RVF. (B) Schematic showing the components of *Voltair$^{PM}$*. C) 15% Native polyacrylamide gel electrophoresis showing the formation of *Voltair$^{PM}$*. Gels were stained with EtBR and imaged in the EtBr, Atto647N as well as RVF channels.

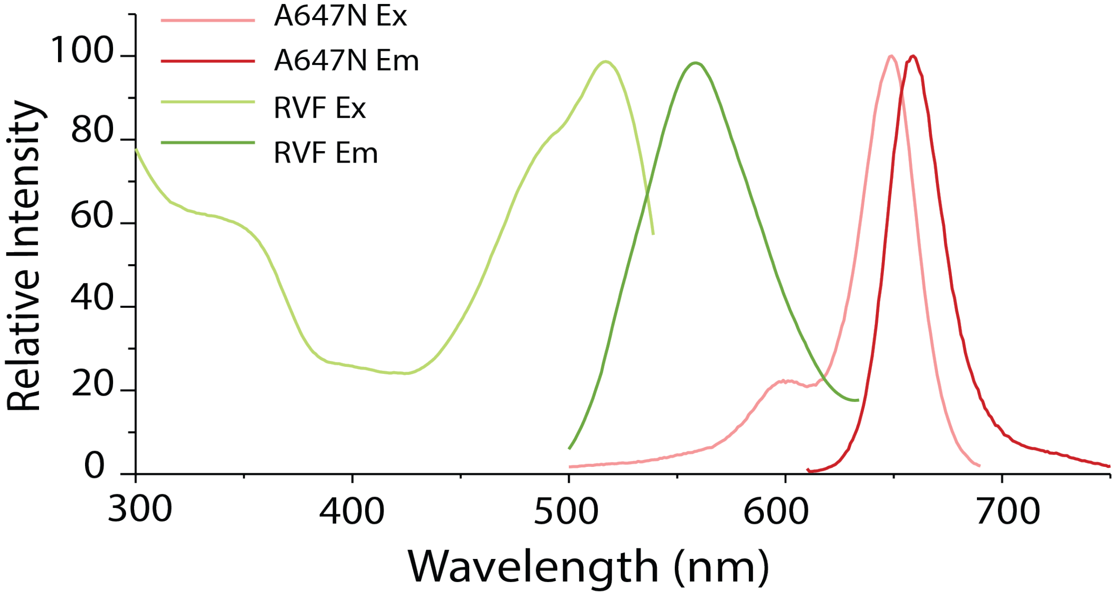


**Fig. S4. Spectral characteristics of *Voltair* probe.** Normalized excitation and emission spectra of RVF (green) and Atto647N (red). The reference dye (Atto647N) was chosen for the minimal overlap of its excitation spectra with the emission spectrum of the sensing (RVF) dye.

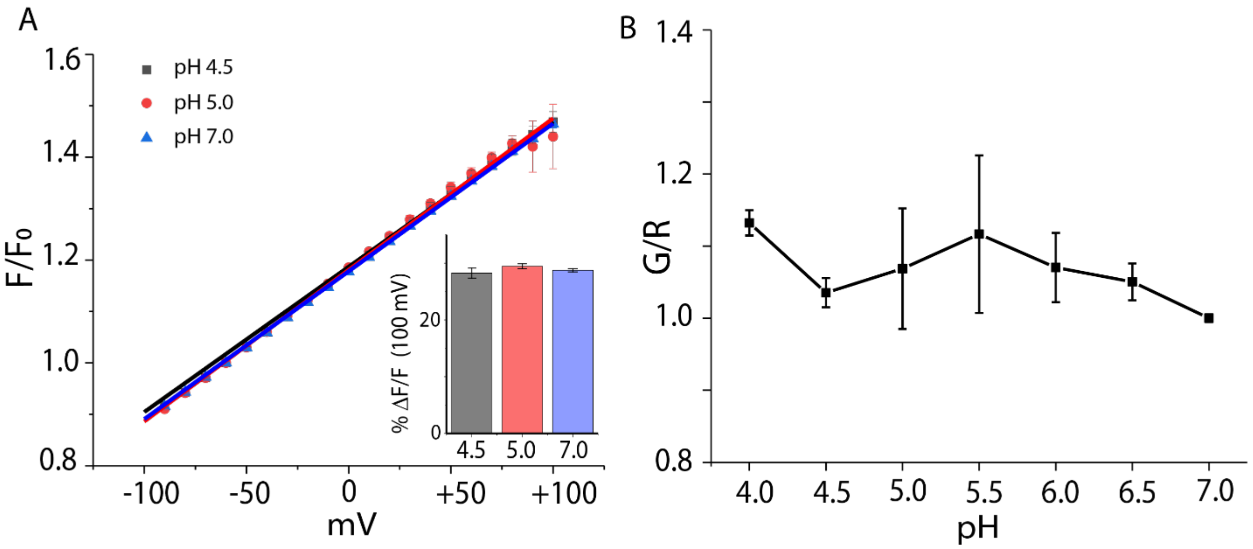


**Fig. S5**. **RVF fluorescence characteristics are insensitive to changes in physiological pH**: A) Voltage sensing characteristics of RVF in the plasma membrane of HEK 293T cells at pH 7, 5 and 4.5 in UB4 buffer. A linear fit to each dataset is plotted. Inset shows the percentage fold change per 100 mV at indicated pH. Error bars indicate s. e. m. of n = 3 experiments. B) Normalized G/R ratios of *Voltair$^{PM}$* recorded in fluorescence spectrometer in UB4 buffers as a function of pH. Error bars indicate S.D. of n = 3 measurements.

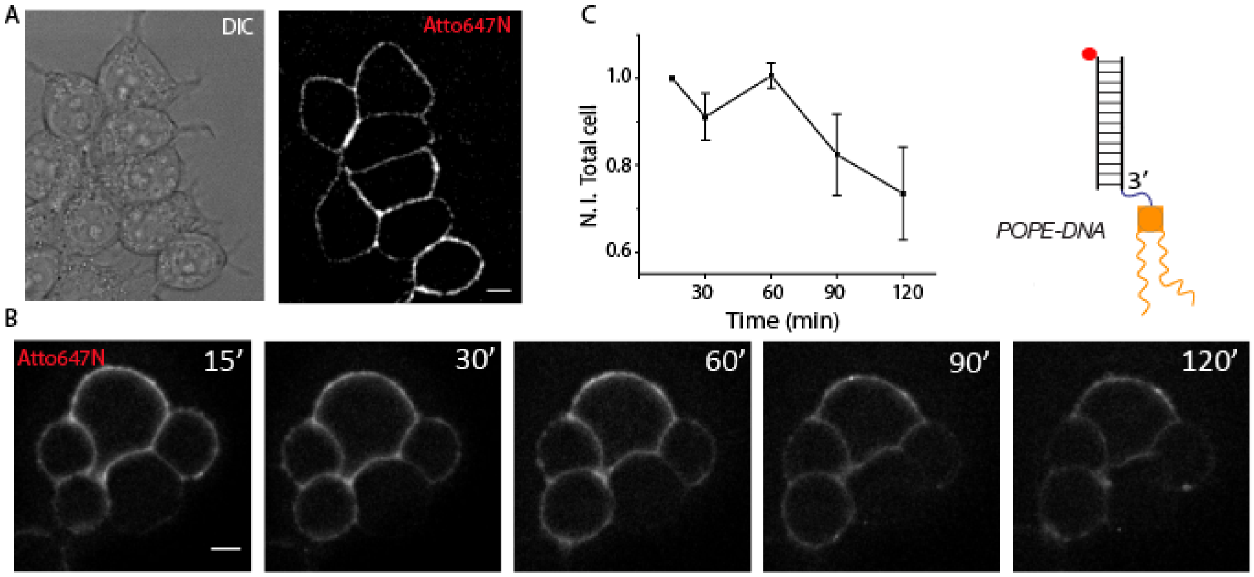


**Fig. S6**. **Plasma membrane labeling with POPE-DNA**. A) POPE-$N_3$ was conjugated to duplex DNA to give POPE-DNA. Addition of POPE-DNA to HEK 293T cells labels the plasma membrane as shown in a representative image of Atto647N. B) Time lapse images of HEK 293T cells labelled with POPE-DNA (500 nM) as a function of time shows a loss of signal only beyond 60 min. Scale = 10 µm C) Normalized total cell intensity (N.I. Total cell) shown in (B) as a function of time. Error bar indicates S.D. of n = 15 cells.

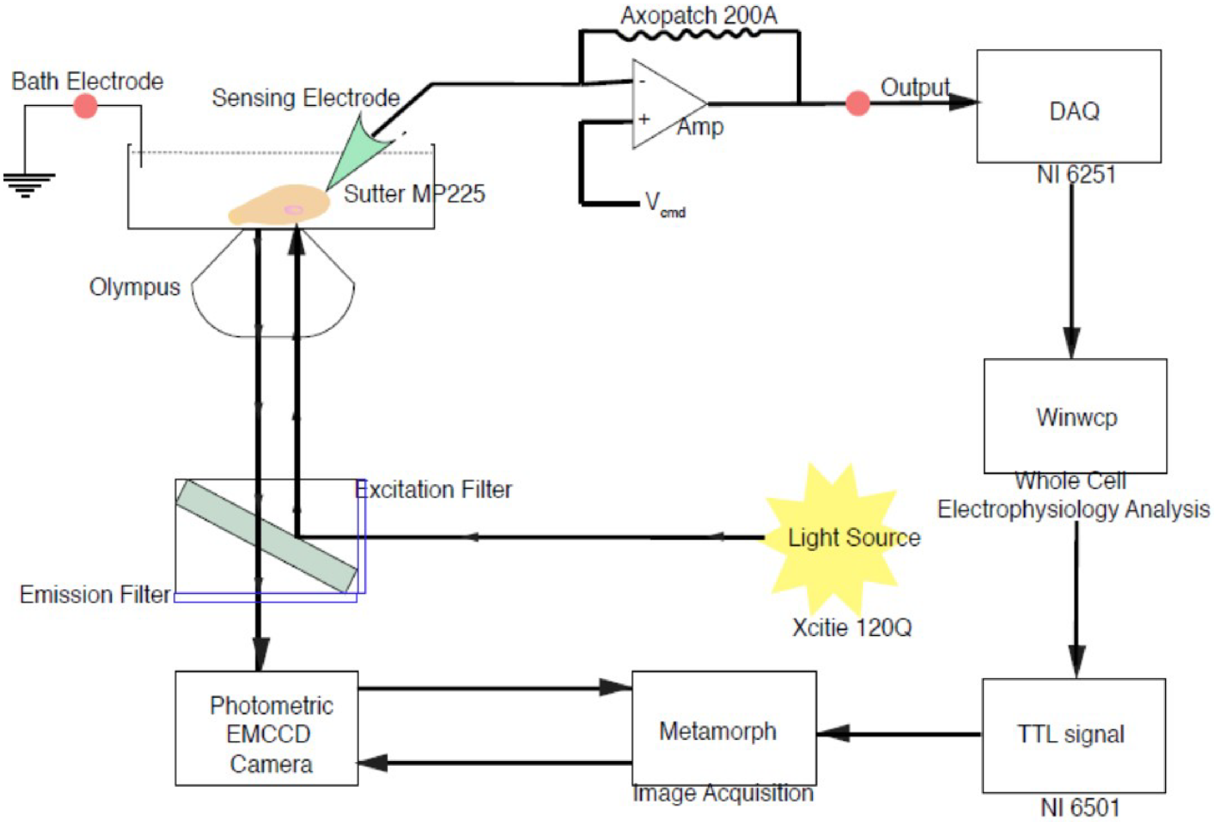


**Fig. S7**. **Optical electrophysiology set up for single cell voltage clamping**. Schematic shows the equipment workflow of electrophysiology measurements. Recordings were performed with an Axopatch 200A amplifier. The signals were digitized with NI-6251 DAQ and recorded using WinWCP software (Strathclyde Electrophysiology Software). The patch pipette was positioned using MP325 motorized manipulator. Image Acquisition software Metamorph premier Ver. 7.8.12.0 was linked to an NI-6501 to enable voltage triggered image acquisition. When applying a voltage pulse across the cell membrane a digital output pulse was generated by WinWCP to trigger imaging. Images are acquired using IX83 widefield microscope, 60X, 1.4 NA, Oil objective.

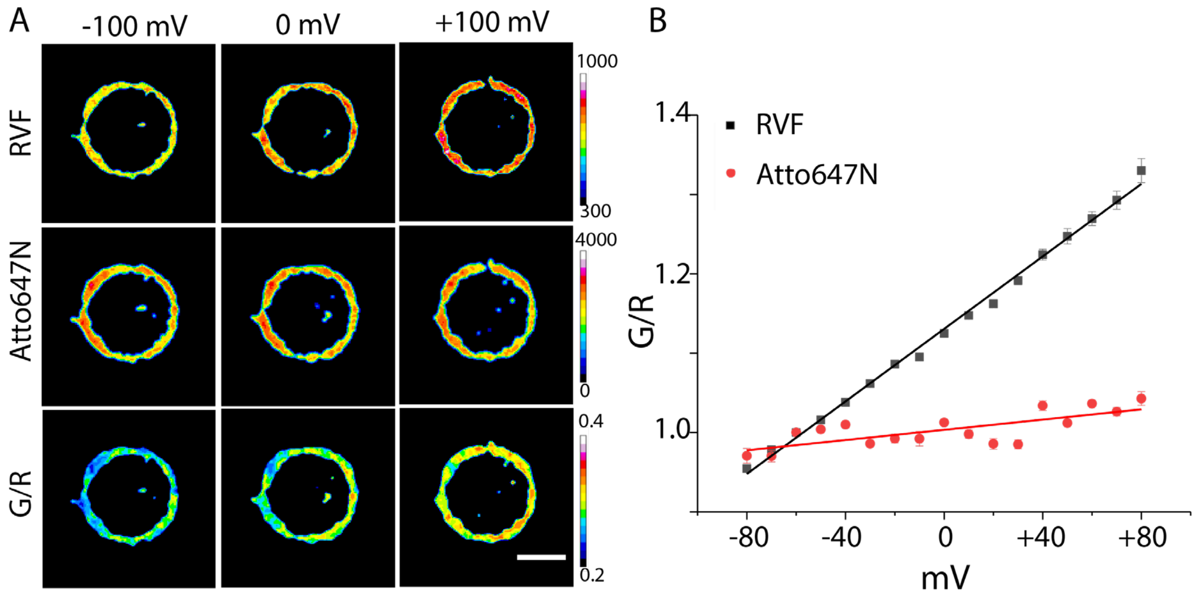


**Fig. S8. Voltage sensitivities of RVF and Atto647N in *Voltair$^{PM}$*:** A) Representative images of *Voltair$^{PM}$* labelled HEK 293T cells voltage clamped from -100mV to +100mV, pseudocolored according to the pixel-wise fluorescence intensities of RVF (G), Atto647N (R) and ratio G/R. Scale bar = 10 µm. B) Normalized RVF (black) and Atto647N (red) intensities of *Voltair$^{PM}$* labelled HEK 293T cells voltage clamped from -80 mV to +80 mV. Error bars indicate s. e. m. of N = 3 experiments.

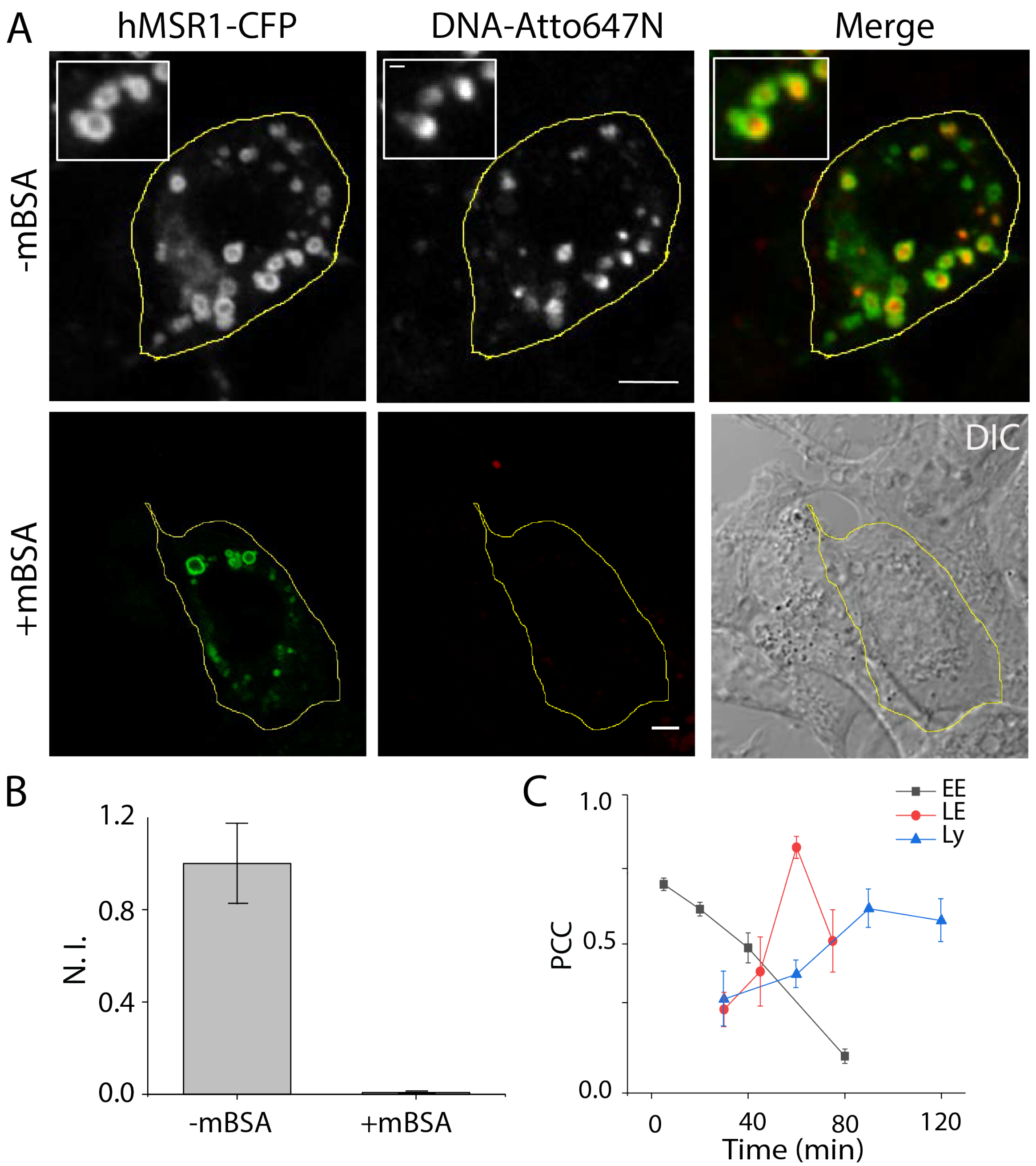


**Fig. S9. *Voltair$^{IM}$* uptake by HEK cells expressing human scavenger receptor (hMSR1).** (A) Upper Panels: *Voltair$^{IM}$* is endocytosed by HEK 293T cells expressing hMSR1-CFP. Colocalization of hMSR1 (CFP channel) with *Voltair$^{IM}$* (Atto647N channel) in punctate structures. Lower panels: Uptake of *Voltair$^{IM}$* is efficiently competed out in the presence of maleylated BSA (mBSA) a good ligand for hMSR1. Scale bar = 10 µm. (B) Normalized avg. intensity of cells shown in (A). (C) Pearson's correlation coefficient (PCC) of colocalization with endocytic markers as a function of *Voltair$^{IM}$* chase times. Error bars indicate the standard deviation for n = 50 cells.

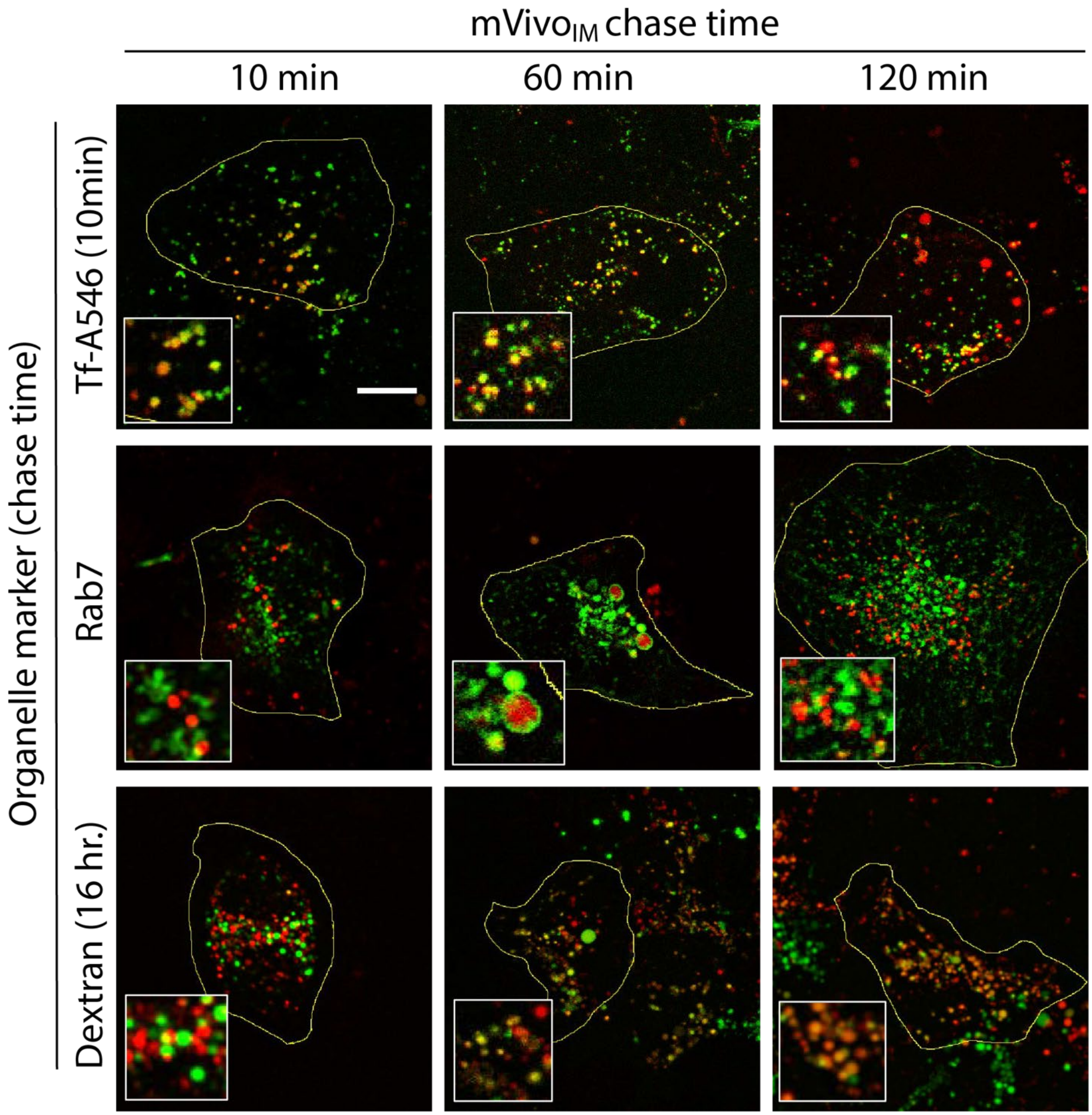


**Fig. S10. Time dependent trafficking of *Voltair$^{IM}$*:** Representative colocalization of singly labeled *Voltair$^{IM}$* (red) and endosomal markers (green) in HEK 293T cells expressing hMSR1. Early endosomes are labeled with 10 min pulse Alexa 546-labeled transferrin (Tf-A546). Late endosomes are labeled with Rab7-mRFP. Lysosomes are labeled by pulsing TMR-Dextran for 1 hour and chasing for 16 hours. *Voltair$^{IM}$* was pulsed for 10 min and chased for indicated time and imaged accordingly. Pearson's correlation coefficients are shown in Fig 2 of the main manuscript. Scale bar = 10 µm.

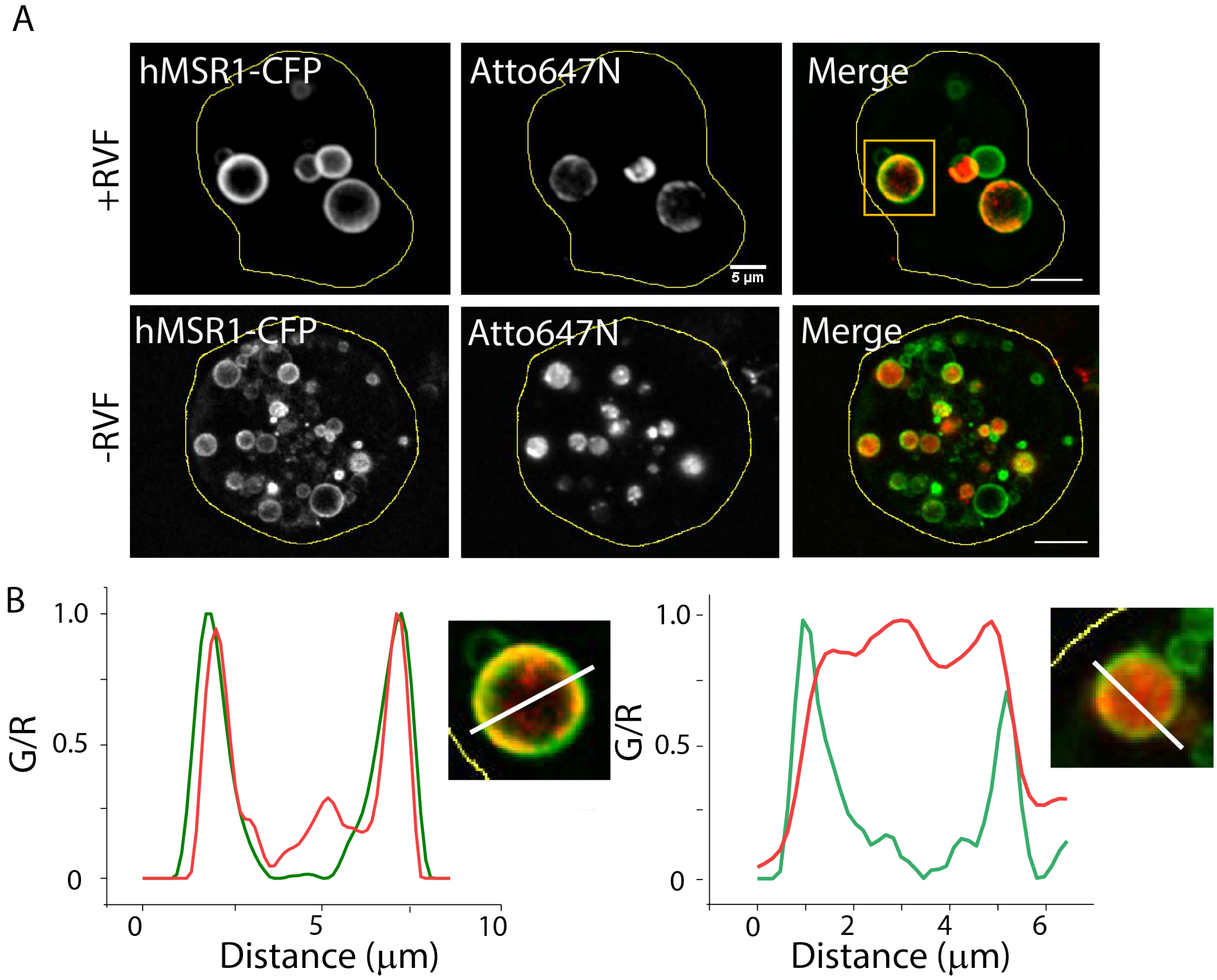


**Fig.S11. *Voltair$^{IM}$* inserts into the membrane of the intracellular organelle because the RVF moiety acts as a lipid anchor**. A) Vacuolin-1 treated HEK 293T cells labeled with *Voltair$^{IM}$* show the DNA probe insert into the lumenal leaflet of lysosomal membranes. When labeled with duplex DNA lacking lipophilic RVF, DNA probe fail to localize at the membrane. B) Line profile of vacuolin-1 treated single lysosome shows the colocalization of DNA probe with hMSR1-CFP present on the endo-lysosomal membrane. Scale = 5µm.

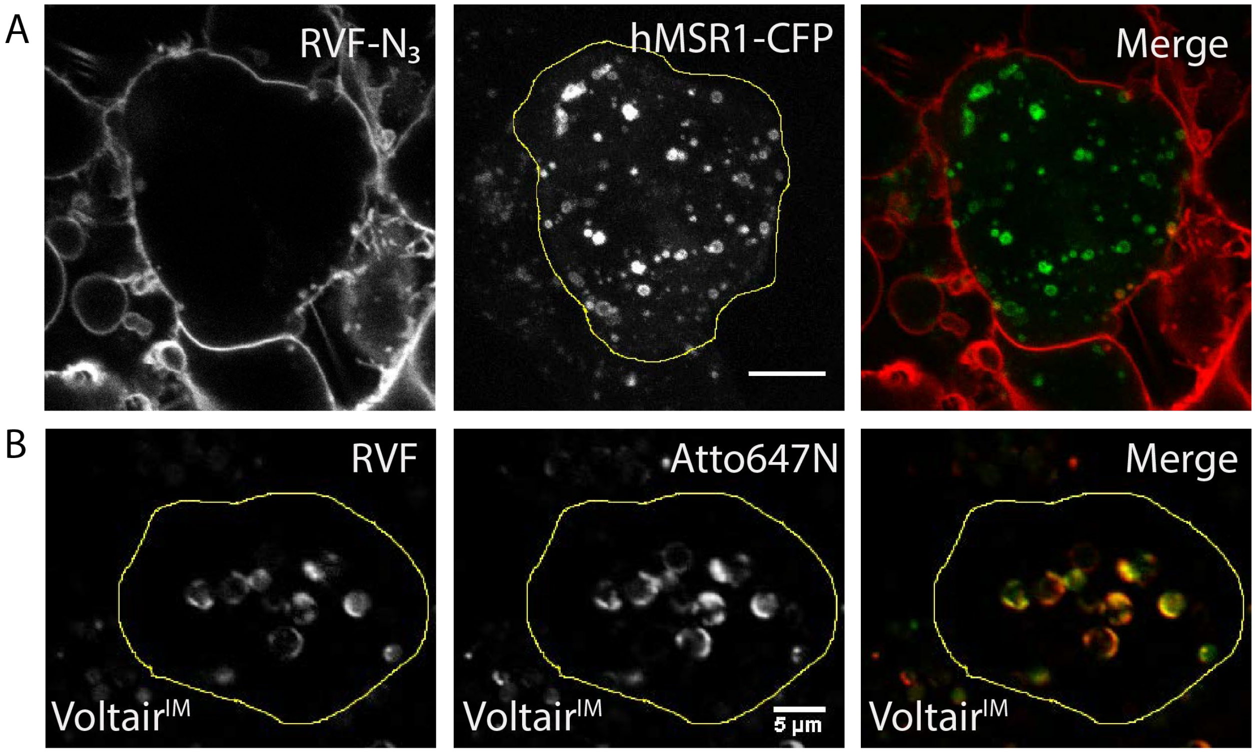


**Fig. S12. Conjugation to the DNA duplex reprograms the affinity of RVF to intracellular membranes**: A) Incubating RVF with hMSR1-CFP expressing HEK 293T cells results in RVF labeling only the plasma membrane. B) RVF is conjugated to duplex DNA to give *Voltair$^{IM}$*. hMSR1 expressing HEK 293T cells incubated with *Voltair$^{IM}$* imaged in the RVF channel and the Atto647N channel (DNA) shows that RVF now labels organellar membranes. Cells were treated with Vacuolin-1 to swell the vesicles in order to confirm tethering to intracellular membranes. Scale: 5 µm.

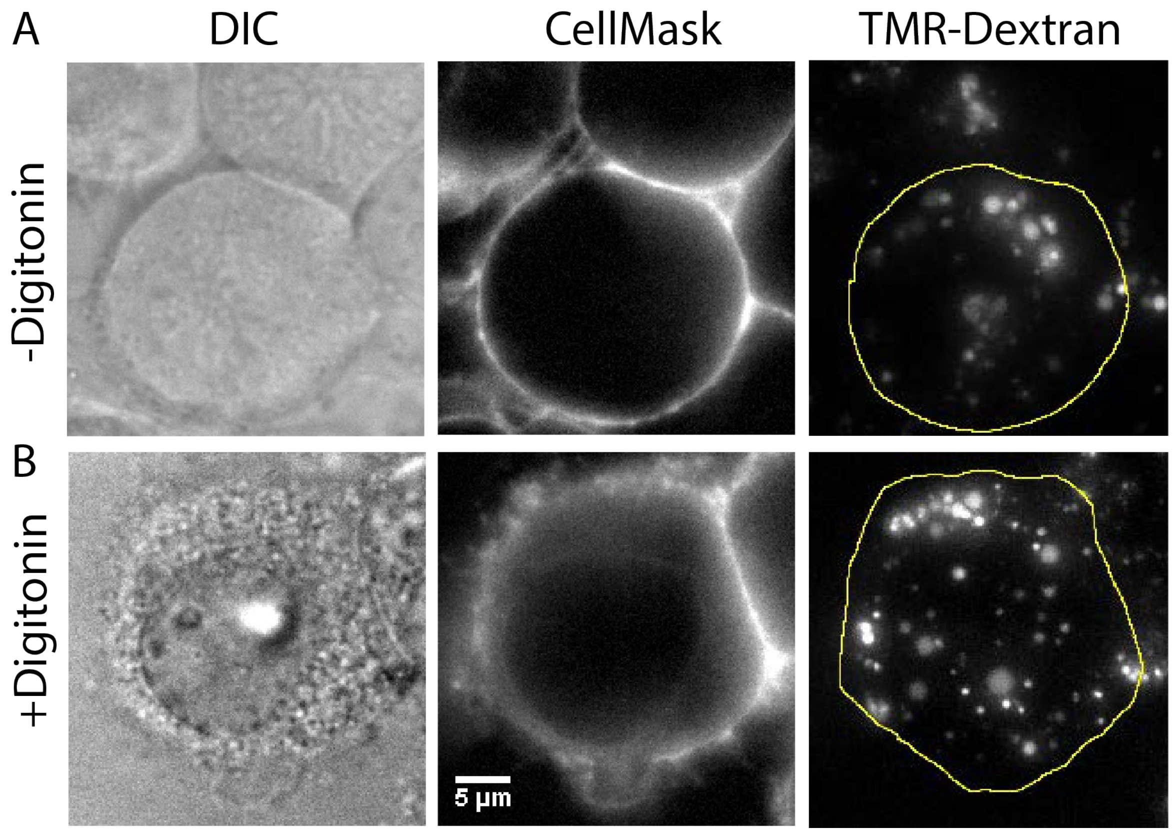


**Fig. S13. Digitonin selectively permeabilizes the plasma membrane**: **(A)** HEK cells incubated with CellMask™ red shows only plasma membrane labeling. Lysosomes labeled with TMR-Dextran are shown for reference. **(B)** HEK cells treated with digitonin and incubated with CellMask™ red now shows labeling of intracellular membranes by CellMask™ red. Note that TMR-Dextran labeled lysosomes are still retained in punctate structures, showing lysosomes are not permeabilized. Scale = 5 µm.

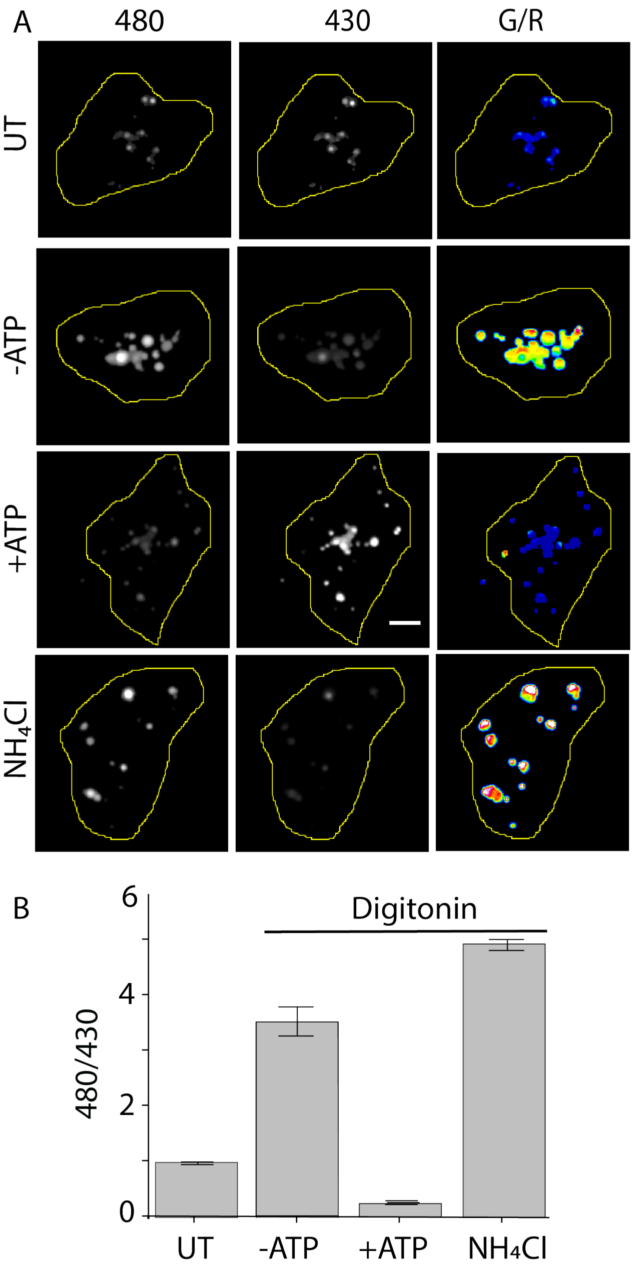


**Fig. S14. Ion-pumps on lysosomes are fully functional post digitonin treatment**: (A) pH of native lysosomes labeled with FITC-dextran imaged using dual excitation method, gives low 480/430 ratio corresponding to native lysosomal pH. (B) Digitonin treatment and incubation in intracellular buffer without ATP increases lysosomal pH as V-ATPases cannot function under ATP-depleted conditions (C) Digitonin treatment in intracellular buffer containing 2 mM ATP decreases lysosomal pH as V-ATPases can now function. (D) Adding $NH_4Cl$ to the buffer in (C) elevates lysosomal pH as expected for functionally intact lysosomes. Scale =10 µm, error bars indicate s. e. m. of n = 2 experiments.

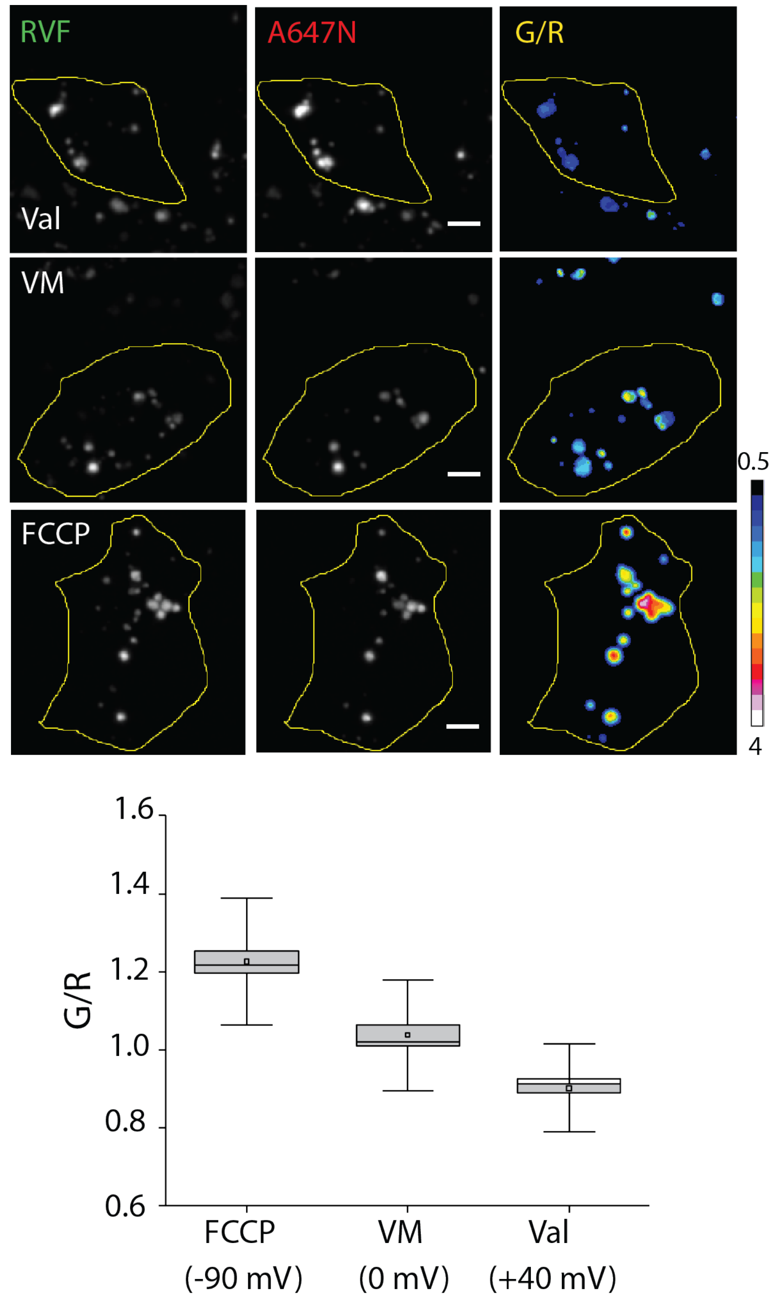


**Fig. S15. Intracellular voltage clamping of *Voltair*$^{IM}$**. (**A**) Fluorescence images of *Voltair*$^{IM}$ labeled lysosomes, voltage clamped using ionophores, in the G and R channels as well as the corresponding pseudocolor G/R images. *Voltair*$^{IM}$ labeled cells were treated with digitonin, incubated with intracellular buffer containing the indicated ionophores to clamp lysosomal membrane potential. (Val – Valinomycin, VM – Valinomycin and Monensin) (**B**) Normalized G/R intensity ratios of each voltage clamped condition in (A). G/R values were normalized to valinomycin and monensin treated cells. Scale = 10 µm, error bars indicate SD of n = 50 lysosomes.

.

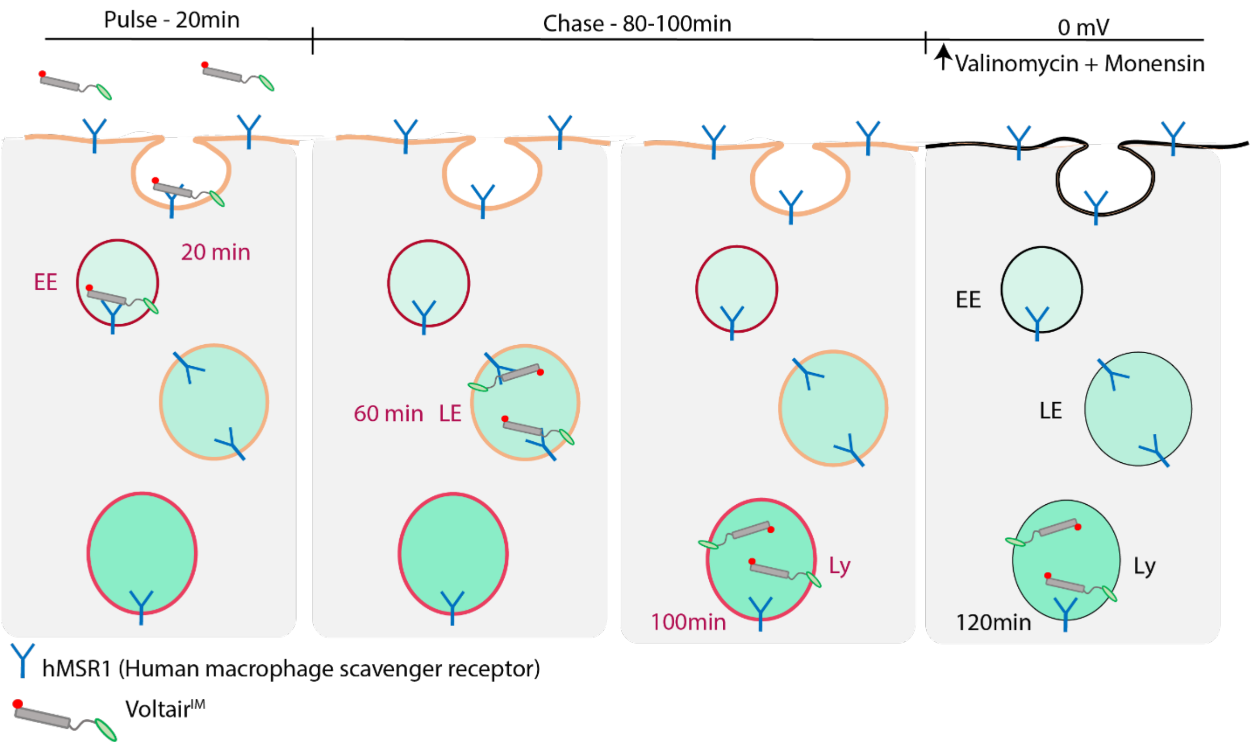


**Fig. S16**. **Schematic representation of pulse-chase protocol for organellar measurements.** HEK cells transfected with hMSR-1 were incubated with *Voltair$^{IM}$*, washed and chased for the indicated chase times to achieve localization in the early endosome (EE), late endosome (LE) and lysosome (Ly) respectively. Images are recorded at indicated chase times. This followed by the addition of a cocktail of valinomycin and monensin to neutralize the membrane potential across all organelles, after which, images are again recorded to obtain the G/R value for 0 mV. G/R values of every organelle are normalized to the G/R value at 0 mV from which organelle membrane potential is quantified.

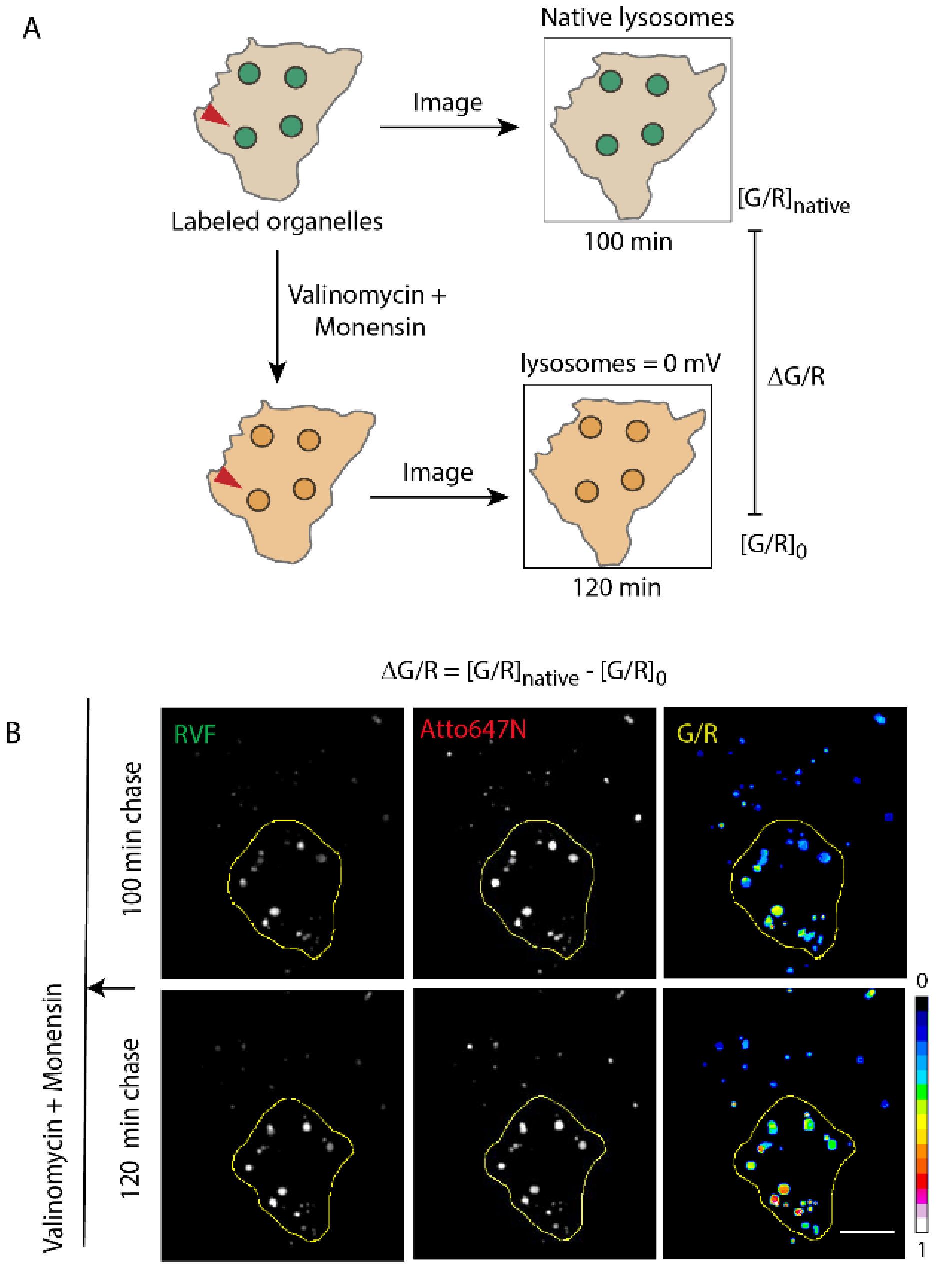


**Fig. S17. Measurement of organelle membrane potential using lysosomes as an example.** (**A**) Schematic of imaging protocol of organelles labeled with *Voltair*$^{IM}$. Native organelles are imaged in G and R channels, neutralized with valinomycin-monensin cocktail and again imaged in the G and R channels. G/R values of native organelle membrane potential are normalized to the G/R values of and neutralized organelle membrane potential (0 mV). (**B**) Representative images of *Voltair*$^{IM}$ labeled lysosomes of the same cell in G and R channels before and after treatment with valinomycin and monensin. Corresponding pseudo-color G/R maps of native and neutralized lysosomes display the membrane potential of each lysosome has increased after treatment.

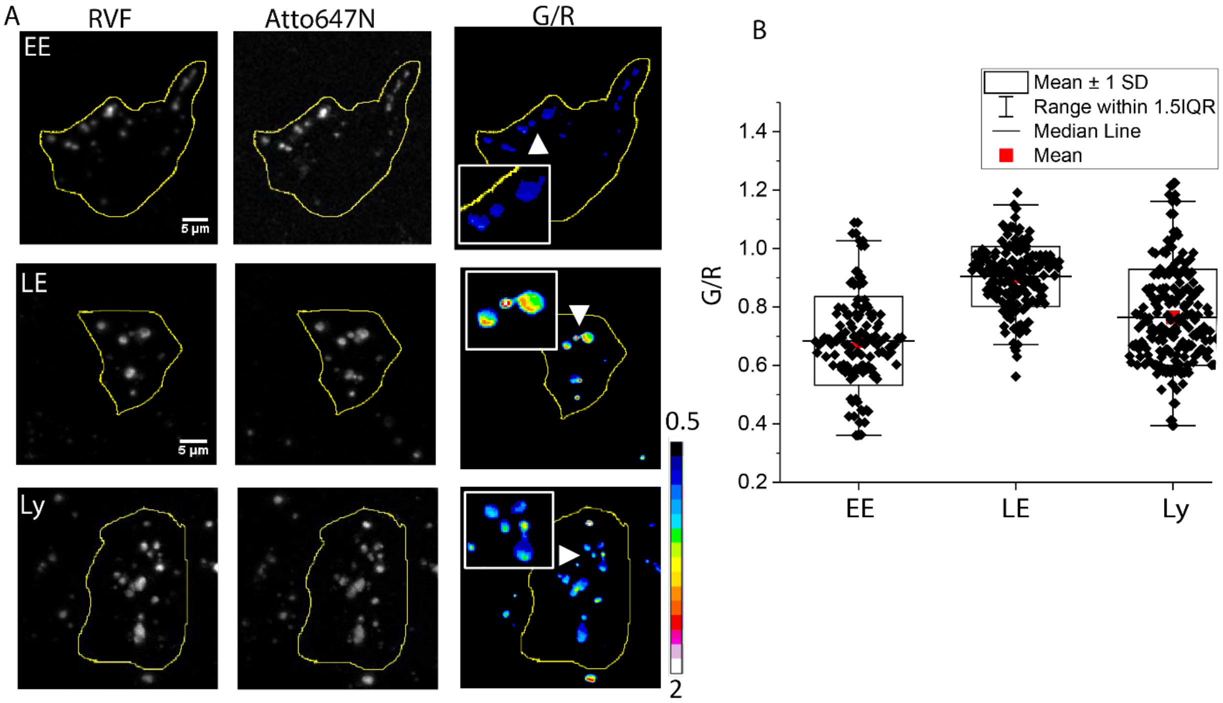


**Fig. S18. Organellar membrane potential measurements**. Representative images of at early endosome, late endosome and lysosomes of HEK 293T cells labelled with *Voltair*$^{IM}$ imaged in RVF, Atto647N and the pseudo-color G/R map. Scatter plot of G/R values of n = 200 organelles. Box represents the standard deviation. Scale = 5 µm.

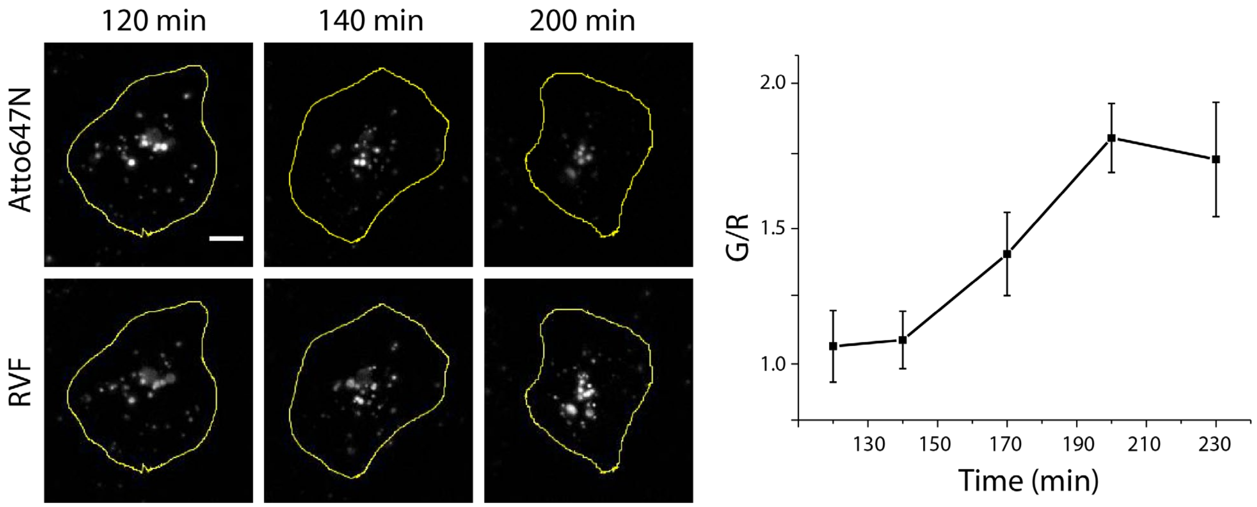


**Fig. S19. Stability of *Voltair*$^{IM}$ in HEK 293T cells:** Stability is evaluated from the fluorescence intensity ratio of non-degradable, membrane impermeable RVF (G) to Atto647N (R), which upon DNA degradation leaks out of the endosome. The ratio of G/R as a function of chase times reveals that *Voltair*$^{IM}$ remains stable up to 140 minutes. Scale = 10 µm, error bars indicate s.e.m. of n = 2 experiments.

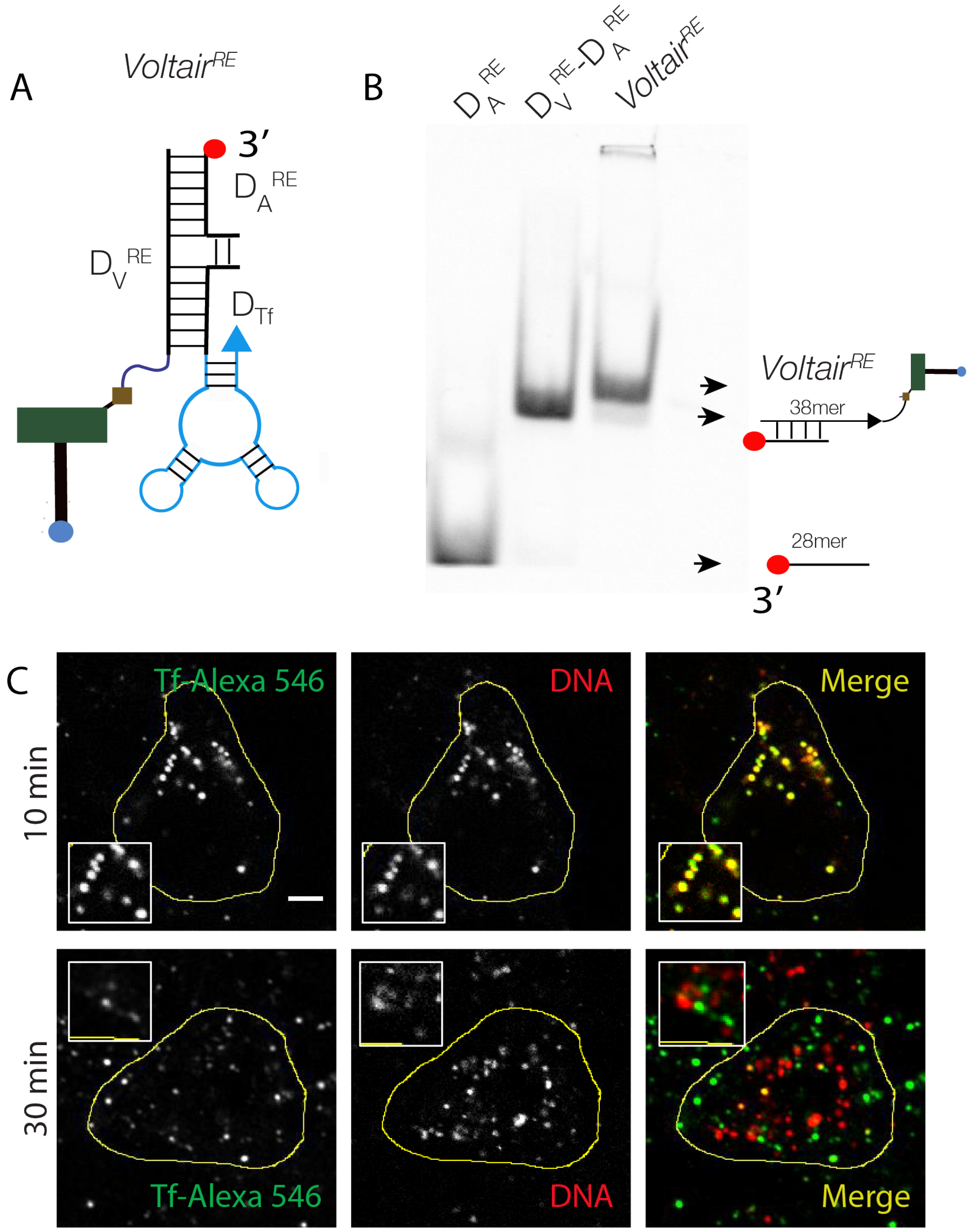


**Fig. S20. Characterization of *Voltair^RE*:** (A) *Voltair^RE* is a trimeric complex comprising of voltage sensing strand $D_V$**^RE**, normalizing strand $D_A$**^RE** and targeting strand $D_{Tf}$. (B) 15% Native polyacrylamide gel electrophoresis imaged in Atto647N channel, showing the formation of *Voltair^RE*. (C) Colocalization of fluorescently labeled transferrin with early endosomes labeled by DNA-Atto647 duplex in 10 mins and 30 mins of chase time (Scale = 5 µm).

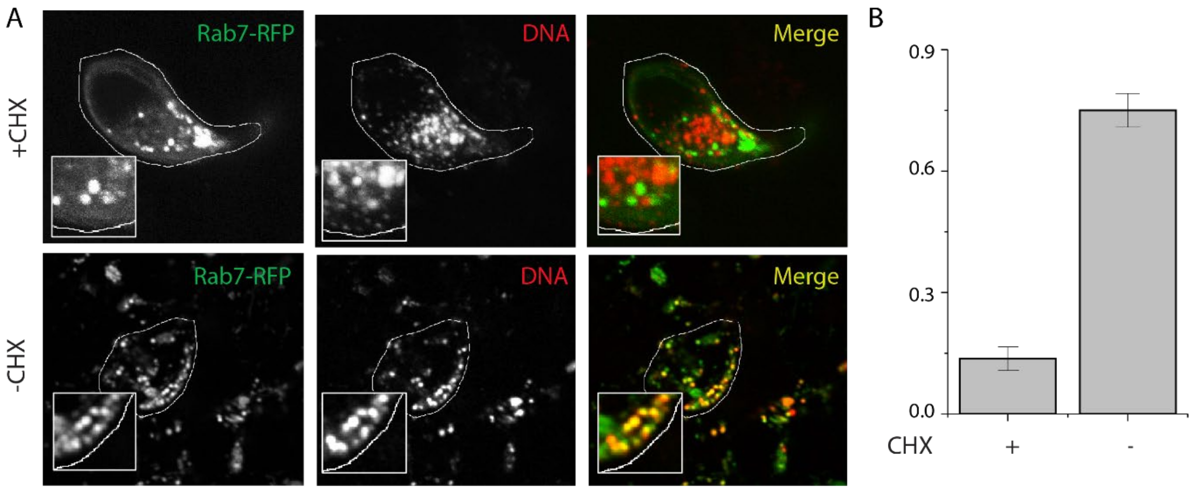


**Fig. S21. Trafficking of *Voltair*$^{TGN}$ in ScFv-Furin expressed cells:** (A) Colocalization of *Voltair*$^{TGN}$ with late endosome marker Rab7-RFP in presence and absence of 125 µg/ mL cycloheximide (CHX). (scale = 5 µm). (B) Pearson's correlation coefficient (PCC) for (A). Error bar represents S.E.M of 15 individual cells.

| S. No | Strand | Sequence (5' to 3') | Comment |
|---|---|---|---|
| 1 | $D_V$ | ATCAACACTGCACACCAGACAGCAAGATCCT ATATATA-ispacer18-DBCON | Sensing stand – conjugated to RVF |
| 2 | $D_T$ | DBCO-TEG-TATATATAGGATCTTGCTGTCT | Targeting strand – conjugated to POPE |
| 3 | $D_A$ | Atto647N – GGTGTGCAGTGTTGAT | Normalizing strand – PM |
| 4 | $D_A$' | TATATATAGGATCTTGCTGTCTGGTGTGCAG TGTTGAT-Atto647N | Normalizing strand – Intracellular Membrane (IM) |
| 5 | $D_V^{RE}$ | DBCO-TEG-ispacer18-TATATATAGGATCTTGCTTCTGTGCCTGCAGTGTTGAT | RE sensing strand – conjugated to RVF |
| 6 | $D_A^{RE}$ | Atto647N-ATCAACACTGCAGGCACAGAGTCTGGTG | RE normalizing stand |
| 7 | $D_{Tf}$ | CACCAGACAGCAAGATCCTATATATAGGGGGA**UC**AA**UCC**AAGGGA**CCC**GGAAA**C**G**UCCCUU**A**C**A**C** **CCC** | RE targeting strand – modified with RNA aptamer. The portion of sequence in red correspond to RNA aptamer against hTfR and bold letters indicate 2' fluoro modified bases. |

**Table S1. Sequences used for *Voltair* probe assemblies.** Strands $D_V$, $D_T$ and $D_A$ form *Voltair$^{PM}$*, $D_V$ and $D_A$' form *Voltair$^{IM}$ and Voltair$^{TGN}$*, $D_V^{RE}$, $D_A^{RE}$ and $D_{Tf}$ forms *Voltair$^{RE}$*.

| Compartments | Membrane potential (mV) |
|---|---|
| EE | 134 ± 17 |
| LE | 40 ± 5 |
| Ly | 100 ± 13 |
| RE | 55 ± 11 |
| TGN | 103 ± 16 |

**Table S2. Membrane potential values for different endocytic organelles.** The error consists of two components, measurement error for G/R and calibration error. Calculation of error is explained above in the Organellar Membrane Potential Measurement section

**Movie S1.**

Pseudo-colored video showing the sensitivity of RVF dye. RVF labeled HEK 293T cells were voltage clamped from -100 mV to +100 mV, at increments of 10 mV in extracellular buffer. The clamped cell is shown by white arrow. Scale bar = 10 µm.

**Movie S2.**

Pseudo-colored video showing the sensitivity of *Voltair$^{PM}$*. Labeled HEK 293T cells were voltage clamped from -100 mV to +100 mV, at increments of 10 mV in extracellular buffer. The RVF channel shows the fluorescence change with respect to membrane potential, whereas Atto647N fluorescence is insensitive to applied voltage increments. The G/R ratio quantitatively shows the change in membrane potential difference. Scale bar = 10 µm.

References:

1. S. Saha, V. Prakash, S. Halder, K. Chakraborty, Y. Krishnan, A pH-independent DNA nanodevice for quantifying chloride transport in organelles of living cells. *Nat. Nanotechnol.* **10**, 645–651 (2015).
2. S. Modi *et al.*, A DNA nanomachine that maps spatial and temporal pH changes inside living cells. *Nat. Nanotechnol.* **4**, 325–330 (2009).
3. E. W. Miller *et al.*, Optically monitoring voltage in neurons by photo-induced electron transfer through molecular wires. *Proc. Natl. Acad. Sci. USA*. **109**, 2114–2119 (2012).
4. N. J. Agard, J. A. Prescher, C. R. Bertozzi, A strain-promoted [3 + 2] azide-alkyne cycloaddition for covalent modification of biomolecules in living systems. *J. Am. Chem. Soc.* **126**, 15046–15047 (2004).
5. D. Moore, D. Dowhan, *Curr. Protoc. Mol. Biol.*, in press, doi:10.1002/0471142727.mb0201as59.
6. B. van Lengerich, R. J. Rawle, S. G. Boxer, Covalent attachment of lipid vesicles to a fluid-supported bilayer allows observation of DNA-mediated vesicle interactions. *Langmuir*. **26**, 8666–8672 (2010).
7. D. Bhatia *et al.*, Icosahedral DNA nanocapsules by modular assembly. *Angew. Chem. Int. Ed. Engl.* **48**, 4134–4137 (2009).
8. A. Vonderheit, A. Helenius, Rab7 associates with early endosomes to mediate sorting and transport of Semliki forest virus to late endosomes. *PLoS Biol.* **3**, e233 (2005).
9. S. Modi, C. Nizak, S. Surana, S. Halder, Y. Krishnan, Two DNA nanomachines map pH changes along intersecting endocytic pathways inside the same cell. *Nat. Nanotechnol.* **8**, 459–467 (2013).
10. C. Grimm, J. Vierock, P. Hegemann, J. Wietek, Whole-cell Patch-clamp Recordings for Electrophysiological Determination of Ion Selectivity in Channelrhodopsins. *J. Vis. Exp.* (2017), doi:10.3791/55497.
11. J. G. Magadán, M. A. Barbieri, R. Mesa, P. D. Stahl, L. S. Mayorga, Rab22a regulates the sorting of transferrin to recycling endosomes. *Mol. Cell. Biol.* **26**, 2595–2614 (2006).
12. J. L. Rosenfeld *et al.*, Lysosome proteins are redistributed during expression of a GTP-hydrolysis-defective rab5a. *J. Cell Sci.* **114**, 4499–4508 (2001).
13. J. van Galen *et al.*, Sphingomyelin homeostasis is required to form functional enzymatic domains at the trans-Golgi network. *J. Cell Biol.* **206**, 609–618 (2014).
14. J. Schindelin *et al.*, Fiji: an open-source platform for biological-image analysis. *Nat. Methods*. **9**, 676–682 (2012).
15. R. G. Johnson, A. Scarpa, Protonmotive force and catecholamine transport in isolated chromaffin granules. *J. Biol. Chem.* **254**, 3750–3760 (1979).
16. R. W. Sabnis, *Handbook of Acid-Base Indicators* (CRC Press, illustrated., 2007).
17. J. E. Whitaker *et al.*, Fluorescent rhodol derivatives: versatile, photostable labels and tracers. *Anal. Biochem.* **207**, 267–279 (1992).
18. M. You *et al.*, DNA probes for monitoring dynamic and transient molecular encounters on live cell membranes. *Nat. Nanotechnol.* **12**, 453–459 (2017).
19. A. E. Vercesi, C. F. Bernardes, M. E. Hoffmann, F. R. Gadelha, R. Docampo, Digitonin permeabilization does not affect mitochondrial function and allows the determination of the mitochondrial membrane potential of Trypanosoma cruzi in situ. *J. Biol. Chem.* **266**, 14431–14434 (1991).

20. M. P. Bradley, D. G. Rayns, I. T. Forrester, Effects of filipin, digitonin, and polymyxin B on plasma membrane of ram spermatozoa—an EM study. *Arch Androl*. **4**, 195–204 (1980).
21. K. Chakraborty, K. Leung, Y. Krishnan, High lumenal chloride in the lysosome is critical for lysosome function. *Elife*. **6**, e28862 (2017).
22. S. Surana, J. M. Bhat, S. P. Koushika, Y. Krishnan, An autonomous DNA nanomachine maps spatiotemporal pH changes in a multicellular living organism. *Nat Commun*. **2**, 340 (2011).
23. Y. Han, M. Li, F. Qiu, M. Zhang, Y.-H. Zhang, Cell-permeable organic fluorescent probes for live-cell long-term super-resolution imaging reveal lysosome-mitochondrion interactions. *Nat Commun*. **8**, 1307 (2017).
